# A dinucleotide tag-based parallel reporter gene assay method

**DOI:** 10.1101/2021.06.29.450267

**Authors:** Naixia Ren, Bo Li, Qingqing Liu, Lele Yang, Xiaodan Liu, Qilai Huang

## Abstract

The causal SNPs leading to increased cancer predisposition mainly function as gene regulatory elements, the evaluation of which largely rely on the parallel reporter gene assay system. However, the common DNA barcode-based parallel reporter gene assay systems are troubled with tag bias, mainly due to the tag nucleotide composition. Here we describe a versatile dinucleotide-tag reporter system (DiR) that enables parallel analysis of regulatory elements with minimized bias based on the next-generation sequencing. The DiR system is also more robust than the classical luciferase assay method, especially in the investigation of moderate-level regulatory elements. We applied DiR-seq assay in the functional evaluation of the prostate cancer risk SNPs in prostate cancer cell lines and disclosed 2, and 6 regulatory SNPs in PC-3 and LNCaP cells. The DiR system has great potential to advance the functional study of risk SNPs that have associations with polygenic diseases.

## Introduction

Genome-wide association studies (GWAS) provide a practical approach for disclosing the genetic variations associated with complex diseases [1]. The GWAS Catalog has recruited over 5000 cancer risk SNPs. More than 90% of these susceptibility SNPs are located outside the protein-coding regions, making it challenging to annotate their functional impacts [2]. Most GWAS SNPs are enriched in the open chromatin regions characterized by the DNase I hypersensitive sites (DHSs) [3–5], and the functional risk SNPs usually alter the chromatin binding of given transcription factors and lead to aberrant target gene expression [6–15]. It indicates that the causal risk SNPs are emerging as potential markers in translational medicine for cancer diagnosis and treatment.

In identifying the functional SNPs from a large number of risk variants, high-throughput reporter gene assay methods are in urgent need. DNA barcodes of 10-20nt at length have been widely used to enable parallel reporter assay of promoters and enhancers [16–19]. However, many factors can affect reporter expression and bring about bias to reporter assay unavoidably, mainly the barcode nucleotide composition. Multiple barcodes such as 90 had to be allocated for each regulatory element to reduce the potential bias effect [18–21]. To explore the position-independent enhancers, the STARR-seq was developed using the self-transcribing active regulatory region as their barcode in NGS sequencing [20,22] and has been applied in several high-throughput screening of risk SNPs [21,23]. However, many of the regulatory SNPs function as distance-dependent proximal regulatory elements [24]. Therefore, we still lack a parallel reporter assay system capable of bias-free gene regulatory function investigation of the risk SNPs, regardless of whether their function is in a distance-independent manner.

Here, we describe a new reporter gene assay system based on dinucleotide barcodes to realize parallel reporter assay with minimized tag bias through the next-generation sequencing. It also enables the reporter expression of individual tags to be quantified through the qPCR method using tag-specific primers. The DiR system exhibited higher consistency, sensitivity, and stability than the present methods, making it more powerful to investigate the subtle regulatory effect from the causal SNPs. DiR-seq application in gene regulatory function assessment of the Prostate cancer (MIM:176807) risk SNPs identified cell-specific regulatory risk SNPs in PC-3 and LNCaP cells. For being accurate, stable, sensitive, and covering general throughput, the DiR system has great potential to advance the study of risk SNPs that have associations with diverse kinds of complex diseases.

## Material and Methods

### Construction of the dinucleotide barcoding system

The 1-450bp DNA region of the luciferase coding sequence was synthesized with G/C content optimized and ‘ATG’ codons removed and then inserted into the pGL3-Promoter vector (E1761, Promega) in place of the original full-length luciferase coding region using ClonExpress®II One Step Cloning Kit (C112-01, Vazyme). The resulted construct was named pDiR001.

Other DiR constructs were prepared through site-directed mutagenesis on the pDiR001 vector using primers listed in **Table S2**. The PCR products containing desired dinucleotide barcodes were digested with DpnI (FD1704, Thermo) and transformed into DH5α competent cells (TSV-A07, Qingke). The next day, the single clones were inoculated into liquid LB medium containing ampicillin and grown at 37 °C for 12-16 hours. The plasmids were extracted using a Plasmid Miniprep Kit (01519KA1, Axygen) and sent for Sanger sequencing (Qingke) for verification.

### Construction of the DiR reporter pool for prostate cancer risk SNPs

We obtained the prostate cancer risk SNP list from the GWAS Catalog (https://www.ebi.ac.uk/gwas/) in 2016, which contained 213 prostate cancer risk SNPs (**Table S1**) at that time. They are tag SNPs reported in the GWAS studies and are significant associated with prostate cancer risk (P-value < 10^-5^). We obtained the 55bp SNP-centered DNA region sequence for both normal and risk alleles from the UCSC genome browser on GRCh38/hg38. The annealed oligos (**Table S3)** were inserted into the DiR vectors between SmaI and BglII sites using T4 DNA Ligase (EL0011, Thermo). Each construct was confirmed correct through Sanger sequencing. The 426 reporter constructs for 213 SNPs were mixed with the DiR-Promoter vector and the DiR-Control vector and subjected to reporter assays in prostate cancer cells.

### Construction of the pNGrR10lib

The CMV enhancer sequence is released from vector pcDNA3.1 and introduced into plasmid pDiR001 between SmaI and HindIII. Then the 10-nucleotides random sequence pool “NNNNNNNNNN” was introduced into the reporter gene region through the PCR method using primers listed in **Table S2**, and the resulted plasmid pool named as p10nt-lib. More than 4 million transformants were obtained in the cloning process to ensure library richness. The purified endotoxin-free plasmid pool was then subjected to cell transfection.

### Cell culture

The LNCaP (ATCC Cat# CRL_1740, RRID: CVCL_1379), PC-3 (ATCC Cat# CRL-1435, RRID: CVCL_0035), and MCF7 (ATCC Cat# HTB-22, RRID: CVCL_0031) cells used in this study were purchased from the American Type Culture Collection (ATCC) and grown in RPMI-1640 (Gibco) supplied with 10% FBS (Gibco) and 1% antibiotics (Penicillin-Streptomycin, Sigma). The Lenti-X 293T cells were purchased from Clontech Laboratories (632180) and maintained in DMEM (Gibco) supplied with 10% FBS (Gibco) and 1% Penicillin-Streptomycin. All the cells were cultured at 37°C with 95% air and 5% CO2 and routinely confirmed to be mycoplasma free using the Myco-Blue Mycoplasma Detector (D101-01, Vazyme). All cell lines used in our study were cultured following the ATCC instructions.

### Cell transfection

Plasmids were transfected cells using Lipofectamine 2000 Reagent (11668-019, Invitrogen), FuGENE HD Transfection Reagent (E2311, Promega), or Polyethylenimine (PEI, 408727-sigma) dependent on cell types. Lipofectamine 2000 was used for LNCaP, and PC-3 cells, FuGENE was used for MCF7, and PEI was used for Lenti-X 293T cells following the manufacturer’s instructions.

Cell transfections were performed at 8-24 hours post cell seeding, depending on the cell density and cell growth status. Generally, a 70-90% confluent cell culture was optimal. The DNA/transfection reagent ratio was 1:3 for Lipofectamine 2000 and FuGENE and 1:1.5 for PEI. The DNA was diluted in Opti-MEM and then added to the diluted transfection reagent. After gently mixing and 10-15min incubation, the DNA complex was added to cells by drops and incubated for 1–2 days at 37°C.

### RNA isolation and reverse transcription

Twenty-four hours post-transfection, cells were washed twice and harvested in 1 × PBS, and Total RNA was extracted from all surviving cells using RNeasy Plus Mini Kit (74136, QIAGEN).

ACCORDING TO THE PRODUCT MANUAL, the RNA was treated with RapidOut DNA Removal Kit (K2981, Thermo Scientific) to remove the trace amount of genomic DNA residue and was then subjected to reverse transcription with High-Capacity cDNA Reverse Transcription Kits (4374967, Applied Biosystems). Briefly, 1.5μg RNA was added into 10μl of 2× RT Master Mix and made up to the final 20μl with nuclease-free water. The reactions were incubated at 25°C for 10 min, followed by 120 min at 37°C, then were inactivated Reverse Transcriptase by heating to 85°C for 5 min. The cDNA products were stored at −20°C or −80°C and ready for qPCR analysis and preparing the NGS sequencing library. For the DiR analysis, the sequence-specific primer BarP6 (CACGATCTGTCCGCACTGCTTGG) was used for reverse transcription, and random primer supplied in the reverse transcription kit was used for reverse transcription for other applications.

### Quantitative PCR

We performed RT-qPCR, using the AceQ qPCR SYBR Green Master Mix (Q111-03, Vazyme) on the thermocyclers Rotor-Gene Q (Qiagen) or LightCycler 96 thermal cycler Instrument (Roche Applied Science) using the program: 95 °C for the 300s of initial denaturation, then 45 cycles of 95°C for 10s, 60°C for 30s, followed by a final extension at 72°C for 10s and with a final program of dissolve curve: 95°C for 15s, 60°C for 30s, and 95°C for 15s. All the qPCR primer pairs were confirmed to have reasonable specificity and amplification efficiency before qPCR assays, and all the qPCR assays were performed in three technical replications. The DiR-qPCR primers are listed in **Table S6.**

### DiR-seq library preparation for Illumina sequencing

The DiR-seq libraries were prepared with two rounds of PCR amplification with cDNA as templates using 2× Phusion Hot Start II High-Fidelity PCR Master Mix (F565L, Thermo Scientific). To adapt the 150 bp paired-end sequencing strategy on the Illumina HiSeq X-TEN platform, we divided the 450bp barcoding region into two amplicons of 271bp and 270bp, respectively, in the first round of PCR. During this step, the binding sites of Illumina sequencing primers were introduced at both ends. In the second-round PCR, adaptors for cluster generation and the index sequences were added. Twenty-four sets of primers tiling the flank sequence of the barcoding region in the first round of PCR, in combination with 12 sequencing indexes introduced in the second-round PCR, will enable up to 288 treatments to be analyzed in parallel in one NGS library. The first-round PCR was performed with 2× Phusion Hot Start II High-Fidelity PCR Master Mix using the program: 98 °C for the 30s of initial denaturation, then 7 cycles of 98°C for 10s, 72°C for 45s and followed by a final extension at 72°C for 5min. The PCR products were purified using 1×VAHTS DNA Clean Beads (N411, Vazyme), eluted in 10μl water, pooled every twenty-four sets of products equally, and then subjected to the second round PCR, which was performed using 2× Phusion HS II HF Master Mix with 1ng template DNA (98 °C for 30 s, 10 cycles of 98 °C for 10 s, 68 °C for 15 s, 72 °C for 30 s, followed by 72 °C for 5min). The products were purified using 1×VAHTS DNA Clean Beads and eluted in 15μl 1×TE buffer. As input control for calculating the expression level, we also subjected the template plasmid pool to NGS library preparation. The purified DiR-seq libraries were subjected to 150 bp paired-end sequencing on the Illumina HiSeq X-TEN platform run by Genewiz, generating about 1 million reads per library. Primers used for DiR-seq library construction are shown in **Table S4**.

### NGS data processing

Illumina sequencing raw data were performed quality control using the software FastP (https://github.com/OpenGene/fastp). It is important to note that the 5’ terminal ‘N’ base should not be removed during the cleaning step. The clean Illumina reads were assembled for the paired reads using the software Panda-seq, and the sub-libraries were then sorted out using the R package ‘ShortRead’. Further, we counted the read number of each dinucleotide barcodes using the R package ‘ShortRead’ for each sub-library and normalized the barcode counts by making each sub-library 1M total reads to eliminate the influence of sequencing depth variation. For each dinucleotide barcode, the expression level was counted by dividing the reads number in cDNA by template DNA. The statistical significance of the expression difference between the two SNP alleles was evaluated with the Two-tailed Student’s t-test. All the SNPs determined to have regulatory functions were listed in **Table S5**.

## Results

### The workflow of the DiR system

The applications of DNA barcodes have enabled parallel reporter assay of regulatory elements and it is popular in the annotation of regulatory element function. The tags with 10-20 nucleotides in length have been widely used in the reporter gene assay [18–21]. Here we designed a new DNA barcoding system using dinucleotide as barcoding tags on a 450bp DNA region. The 450bp barcoding region was generated based on the firefly luciferase coding sequence of the pGL3-Promoter vector by removing the start codon and optimizing nucleotide composition with G/C content (**Supplemental Sequence**). The interested regulatory element can be inserted upstream of the SV40 promoter, and the reporter expression level of the regulatory element can be determined by next-generation sequencing technology and the qPCR method using tag-specific primers after plasmids pool transfection (**Fig. 1**). The DiR reporter system makes report analysis convenient, simple, and efficient.

**Figure 1.**
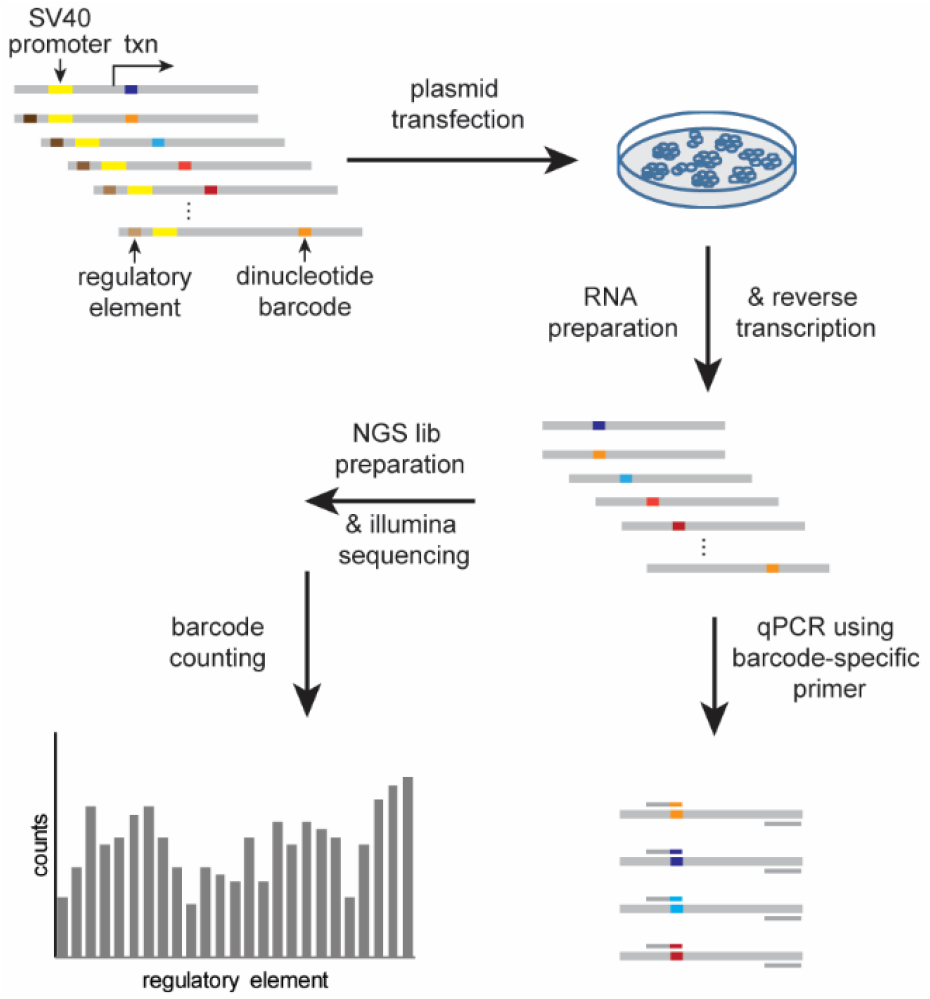
Flowchart of the DiR system for reporter gene assay. The DiR plasmid library was developed based on the pGL3-promoter vector by optimizing the luciferase coding sequence. SV40 promoter shown in yellow, and the dinucleotide barcodes designed on the luciferase sequence shown in colors, and regulatory element inserted upstream of the SV40 promoter.

### Design the DiR system

We previously reported that using primers with the tag-specific nucleotide placed at its 3’ end could achieve sequence-specific PCR quantification, and the nucleotide type of tag could significantly affect the PCR specificity [25]. To determine the best dinucleotide barcoding rule, we constructed 16 plasmids containing all the 16 kinds of dinucleotides using the coding sequence of the HOXB13 gene (**Fig. 2A**) and systematically investigated the specificity of the 16 dinucleotides in qPCR amplification. The qPCR results showed that the dinucleotide barcode-specific primers had extremely high capabilities of up to 2^21^ on average for all the sixteen dinucleotide tags in discriminating against other barcodes that differ at both nucleotide positions (**Fig. 2B**). Based on the tagging rule that the nucleotides should be mutually different at both positions, the sixteen dinucleotides can be sub-grouped into twenty-four blocks according to the equation (16×9×4×1)/4! =24 (**Fig. 2C**). We found that the tag-specific primers from different blocks exhibited differential ability to discriminate between on-target and off-target dinucleotide tags. Considering that the higher selectivity of the tag-specific primers will enable more accurate quantification of each dinucleotide tag in the DiR-qPCR assay, we chose the top nine blocks covering all the 16 dinucleotide tags and in which the dinucleotide tags exhibiting less nonspecific amplification strength from off-target templates for subsequent dinucleotide design. The sixteen dinucleotide tags in the selected nine blocks are highlighted in blue in **Fig. 2C**. Using the same barcoding rule, we designed 628 dinucleotide tags on the 450bp DNA region and generated 628 reporter constructs through site-directed mutagenesis.

**Figure 2.**
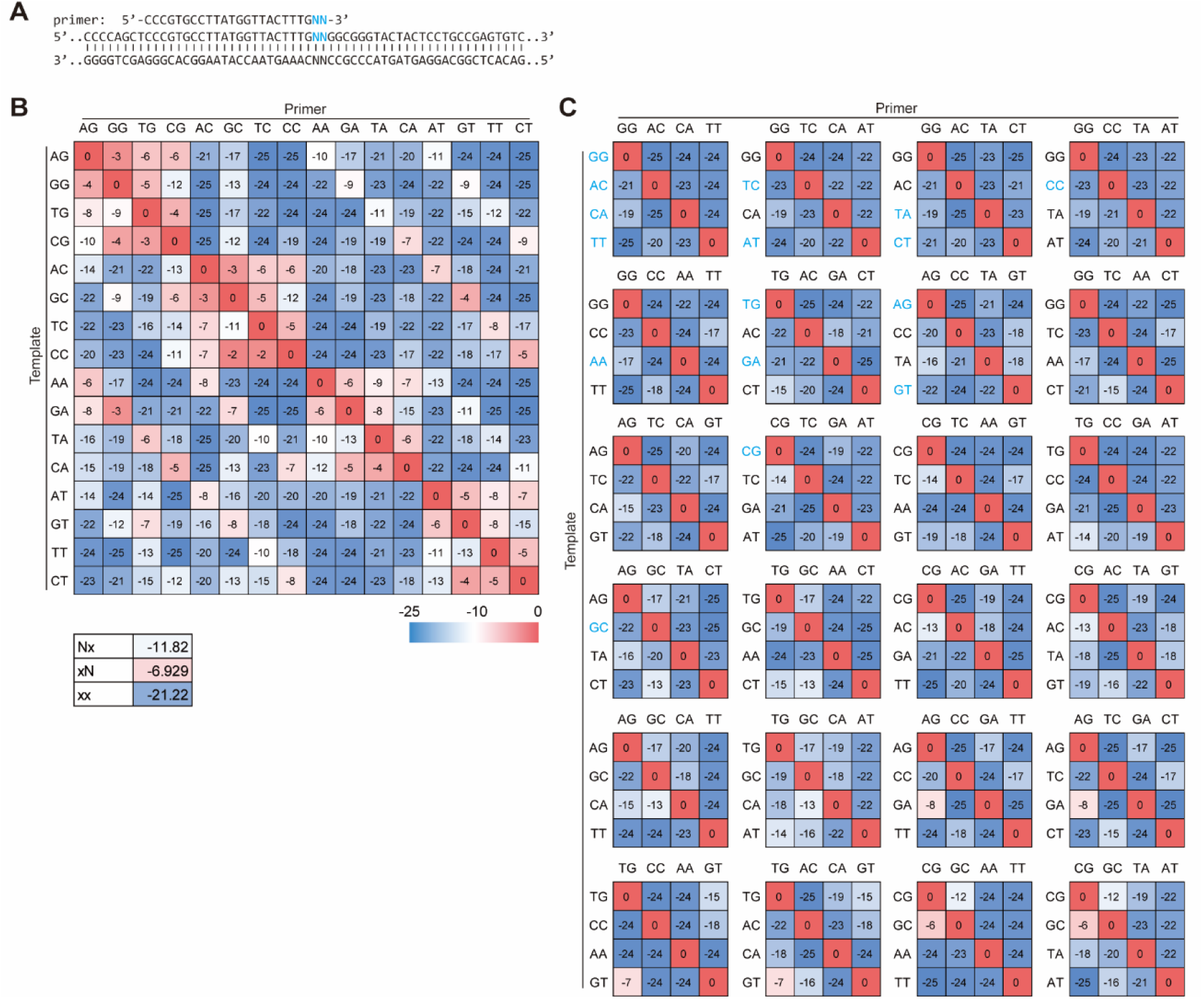
Design the dinucleotide tags. (**A**) Illustration of primer and template used for determining the dinucleotide barcoding rules. (**B**) Relative PCR amplification signal of each dinucleotide-specific primer to the matched and mismatched templates, represented as −ΔCt. The ‘Nx’ represents all the primer-template pairs with a mismatch at the primer 3’ last base. The ‘xN’ and ‘xx’ corresponded to mismatch at the second last base and the last two bases, respectively. The larger absolute values mean lower nonspecific amplifications. (**C**) All the 24 dinucleotide barcoding blocks ranked by their amplification specificity, from left to right and top to bottom. Dinucleotides exhibiting less amplification signal from mismatching templates were selected for barcoding and highlighted in blue.

### The DiR has reduced tag bias

It has been widely recognized that the common tags with 10-20 nucleotides in length could introduce obvious bias on reporter expression levels [18–21]. We then characterized the blank dinucleotide reporter system with comparison to an NGS-based expression assay of a random 10-nucleotide barcoding library encoded in the luciferase coding region. Compared to the random 10-nucleotide barcodes that had a coefficient of variation (CV) value of 88.9% in LNCaP, the expression level of our dinucleotide tags exhibited higher consistency with CV values of 10.2% and 8.5% in LNCaP and MCF7 cells, respectively (**Fig. 3A**). We found that the tag expression levels were affected dramatically by the nucleotide composition in the random 10-nucleotide barcoding library. Notably, the tag expression levels strongly correlated with its GC content (Spearman correlation (r)=0.96, P=7.32e-06; Pearson correlation (r)=0.67, P=0.036) (**Fig. 3B-D**). For the DiR system, only two or four nucleotides differ between each reporter construct on the whole 450bp DNA region, which makes it less affected by the GC composition.

**Figure 3.**
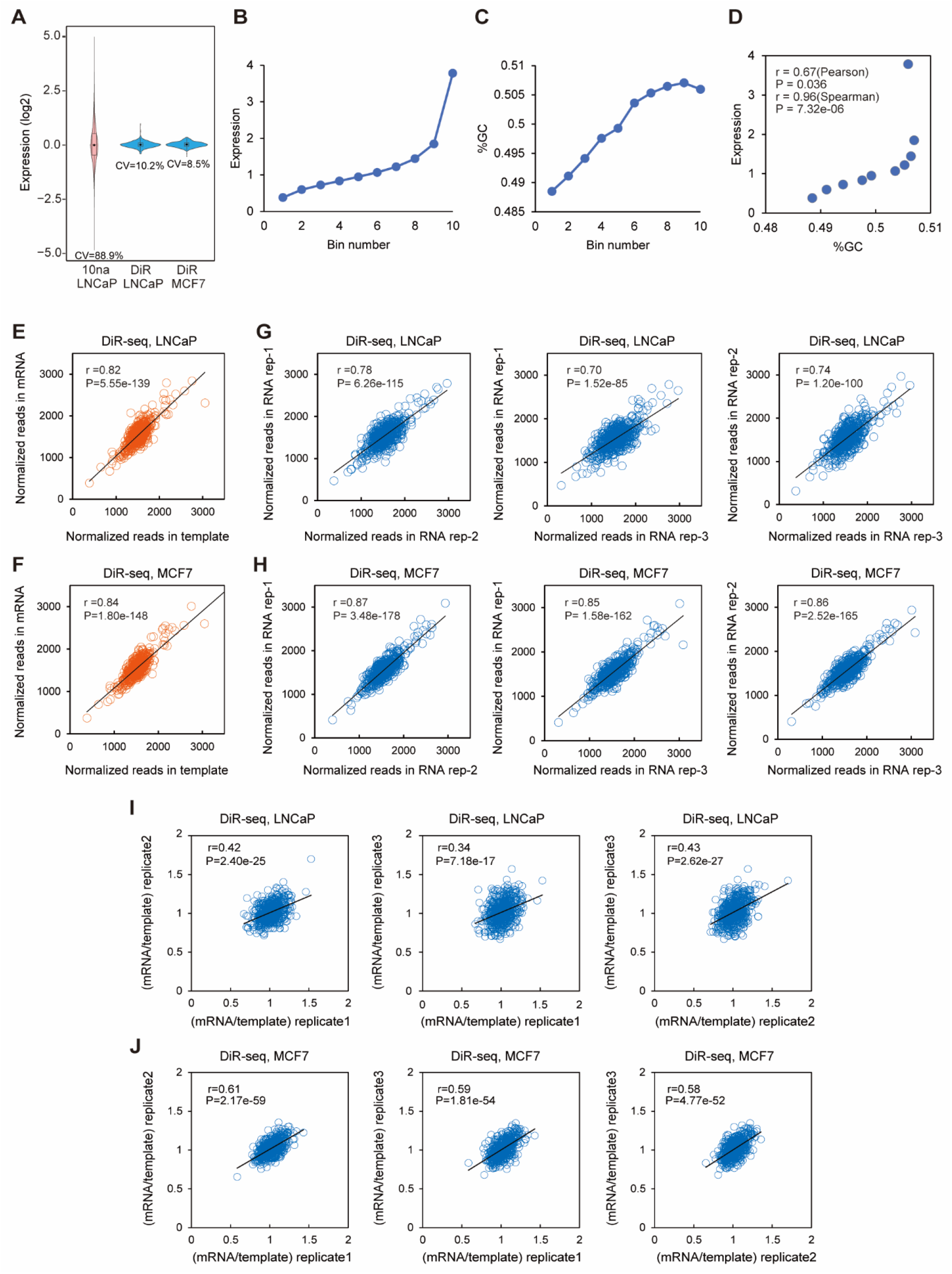
The DiR system has a low bias in the reported assay. (**A**) Violin box plot of the expression level of the 10-nucleotide barcodes in LNCaP cells and dinucleotide tags in LNCaP and MCF7 cells. CV, coefficient of variation. (**B**) The average expression level of the 10-nucleotide barcodes that allocated into ten bins according to their expression level. (**C**) GC content of the 10-nucleotide barcodes in each bin. (**D**) Correlation between tag expression level and GC content percentage of barcodes at ten bins. Correlation coefficient values (r) and P values were calculated with Pearson correlation and Spearman correlation. (**E, F**) Scatter plot of tag reads of the blank DiR-seq in mRNA and template DNA in LNCaP (E) and MCF7 (F) cells, respectively. Correlation coefficient values (r) and P values were calculated with Pearson correlation. (**G, H**) Correlation of tag reads between individual biological replicates in LNCaP(G) and MCF7(H) cells. Correlation coefficient values (r) and P values were calculated with Pearson correlation. (**I, J**) Scatter plot of tag reads ratio of the blank DiR-seq between mRNA and template DNA in LNCaP (I) and MCF7 (J) cells, respectively. Correlation coefficient values (r) and P values were calculated with Pearson correlation.

Moreover, the normalized barcode reads in mRNA and template plasmid library exhibited high consistency for DiR-seq analysis in both LNCaP and PC-3 cell lines (**Fig. 3E, F**), and the normalized reads in mRNA were also consistent between biological replicates (**Fig. 3G, H**). Notably, the template-normalized tag expression levels were much uniform between tags for all the three biological replicates in LNCaP and MCF7 cells. The tag expression level of mRNA divided by template in blank DiR-seq exhibited a moderate correlation between biological replications in both LNCaP and MCF7 cells. (**Fig. 3I, J**). It indicates that even though minimized, the DiR-seq analysis might still have a low-level tag bias. Briefly, we developed a dinucleotide-based reporter gene assay system that enables both parallel and general throughput assay of regulatory elements at high accuracy.

### Comparison of DiR with luciferase reporter assay

We then compared the DiR system with the traditional luciferase assay method. The DiR system does not involve the protein translation process during reporter gene assay because the start codon ‘ATG’ has been removed when designing the dinucleotide barcodes. It allows the DiR system to bring about less stress to the host cells and enable a more robust reporter gene expression than the traditional luciferase reporter assay. In luciferase assay, the normalized luciferase activity of pGL3-Basic, pGL3-Promoter, and pGL3-Control vectors was 1, 11, and 646, respectively (**Fig. 4A**). We observed a similar luciferase expression pattern at the RNA level determined by RT-qPCR analysis (**Fig. 4B**). Notably, in the DiR-qPCR assay, reporter gene expression exhibited a different pattern, for which the relative expression levels of the DiR-Basic, DiR-Promoter, and DiR-Control were 1, 32, and 454, respectively (**Fig. 4C**). The DiR-Basic has no promoter or enhancer element for driving reporter gene expression, the DiR-Promoter has one SV40 promoter upstream of the reporter gene, and the DiR-Control has the CMV enhancer/CMV promoter upstream. The DiR-Promoter generated a significantly higher reporter expression level than the luciferase assay of pGL3-Promoter, but the DiR-Control exhibited a slightly lower level than the pGL3-Control vector. Considering the two control constructs have different promoters and enhancers, and the enhancers were placed in different places relative to the reporter gene, the differential reporter gene expression from the two control vectors might mainly reflect the promoter/enhancer strength and their relative locations.

**Figure 4.**
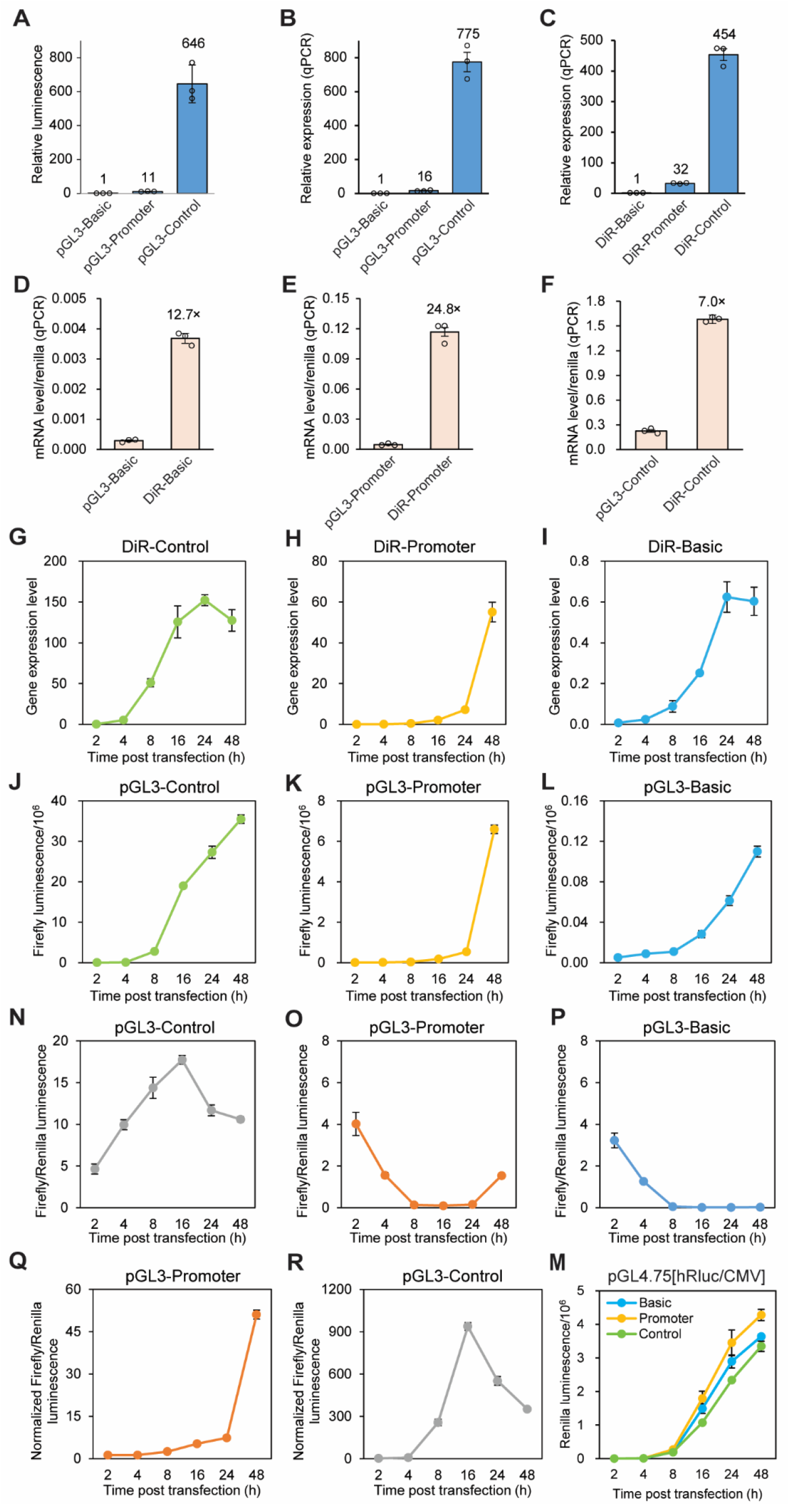
The character of DiR and comparison with luciferase system. (**A**) Relative luciferase activity of pGL3-basic, pGL3-promoter, and pGL3-control in Lenti-X 293T cells at 24h post-transfection. (**B**) The luciferase mRNA level of the three pGL3 luciferase reporter vectors in Lenti-X 293T cells determined through RT-qPCR assay. (**C**) Relative tag expression level from the DiR-Basic, DiR-Promoter, and DiR-Control in Lenti-X 293T cells at 24h post-transfection. (**D-F**) The mRNA level of DiR tags and firefly luciferase normalized to the *Renilla* luciferase in Lenti-X 293T cells for -Basic (D), -Promoter (E), and -Control (F) vectors. (**G-I**) The time-course expression level of tags in Lenti-X 293T of DiR-Control (G), DiR-Promoter (H), and DiR-Basic (I) vectors at 2-48h post-transfection. (**J-L**) The time-course firefly luciferase luminescence of pGL3-Control (J), pGL3-Promoter (K), and pGL3-Basic (L) vectors at 2-48h post-transfection in Lenti-X 293T cells. (**M**) The time-course *Renilla* luciferase activity in Lenti-X 293T cells at 2-48h post-transfection. (**N-P**) The time-course firefly luciferase activity normalized to the *Renilla* luciferase luminescence of pGL3-Control (N), pGL3-Promoter (O), and pGL3-Basic (P) vectors at 2-48h post-transfection in Lenti-X 293T cells. (**Q-R**) The time-course firefly/*Renilla* luciferase luminescence of pGL3-promoter (Q) and pGL3-control (R) vectors calibrated to the value of pGL3-basic at each time point. All the data shown in figure 4 are Mean ± SEM of three independent experiments.

To further compare the dinucleotide tag and luciferase gene directly for their mRNA expression levels, we co-transfected the three DiR plasmids together with individual luciferase report plasmids into Lenti-X 293T cells. We evaluated the reporter gene expression levels with the qPCR method. As expected, the DiR tags showed significantly higher expression levels than the corresponding luciferase vectors, with times of 12.7×, 24.8×, and 7.0× for the DiR-Basic, DiR-Promoter, and DiR-Control vectors, respectively (**Fig. 4D-F**). It suggests that the DiR system allows relatively more robust reporter expression than the traditional luciferase assay, especially for the moderate regulatory elements. It will make the DiR system more effective in screening the risk SNPs that usually drive gene expression at modest levels.

Further, we compared the DiR system with luciferase assay in a time-course manner. The DiR-qPCR analysis showed that the tag expression of DiR-Control, DiR-Promoter, and DiR-Basic all could be detected as early as 2 hours post-transfection and reached the highest point at 24-48h post-transfection (**Fig. 4G-I**). In the luciferase activity assay, the luminescence intensity of pGL3-Control, pGL3-Promoter, and pGL3-Basic displayed similar time-course patterns to the DiR system (**Fig. 4J-L**). However, unlike the DiR system that can deliver all the reporter constructs in one transfection, the luciferase reporter assay system has to transfect individual reporter constructs separately. To eliminate the transfection efficiency differences between constructs, including an internal control reporter such as *Renilla* luciferase or β-galactosidase vector is usually obligatory in each transfection. The common *Renilla* luciferase reporter vector GL4.75[hRluc/CMV] expresses *Renilla* luciferase driven by the strong CMV enhancer/promoter. After transfection, the *Renilla* luciferase activity increased gradually over time with a similar pattern to the pGL3-Control vector (**Fig. 4M**), which caused the normalized luciferase luminescence to deviate dramatically from the real reporter expression levels (**Fig. 4N-P**). For pGL3-Control, the normalized luminescence reached the top point at 16h post-transfection and then declined gradually (**Fig. 4N**). However, for pGL3-Promoter and pGL3-Basic, the normalized luminescence even descended during the beginning 24 hours (**Fig. 4O, P**). It may bring about puzzling results in most reporter gene assay applications, especially those demanding time-course observations. If further calibrated using luminescence of pGL3-Basic at each time point, the reporter expression levels of pGL3-Promoter looked normal (**Fig. 4Q**), but not for the pGL3-Control (**Fig. 4R**). These results demonstrate that the DiR system is more straightforward, reliable, and understandable than the traditional luciferase reporter assay.

### DiR assay discovers regulatory SNPs

The majority of the cancer risk SNPs function as potential gene regulatory elements [5,9,10]. Notably, up to 57.1% of GWAS SNPs locate in the DHSs, which indicates that most of the GWAS SNPs have potential regulatory functions themselves [5]. Hence, we then applied the DiR system to evaluate the 213 prostate cancer risk SNPs with both alleles from the GWAS catalog for their gene regulatory function (**Table S1)**. We cloned the 55bp DNA regions with individual alleles in the middle and inserted them into the DiR vectors right upstream of the basic SV40 promoter. The typical transcription factors only occupy 6-12bp DNA sequences [26], and short DNA stretches bearing transcription factor binding sites were widely used in the reporter gene assay [27]. Therefore, assessing the gene regulatory function of SNPs using the 55bp DNA regions is reasonable, even though we might not observe the full activity of the potential herein enhancer element.

Our DiR-seq analysis in PC-3 and LNCaP cells showed that the tag expression levels had high consistency between individual replicates (**Fig. 5A, B**), and some SNP sites exhibited elevated tag expression levels (**Fig. 5C, D**). Because the two alleles of the functional SNPs are supposed to drive gene expression differentially, we picked SNPs based on the ratio of reporter expression level from the risk and normal alleles (**Fig. 5E, F**). In the PC-3 prostate cancer cell line, 5 SNPs exhibited decreased gene expression for the risk alleles compared to the normal alleles (P<0.05), and 4 SNPs showed increased expression to the contrary (P<0.05) (**Fig. 5E**). Correspondingly, in the LNCaP cell line, 5 SNPs manifested downregulated reporter activity for the risk alleles, and 5 SNPs had upregulated reporter activity (**Fig. 5F**). Considering both the allele preference and the reporter level, we highlighted two functional SNPs, rs5945572 and rs10187424 in PC-3 cells and six SNPs, rs10795917, rs7153648, rs17765344, rs1327301, rs10187424, and rs887391 in LNCaP cells in Figure. 5E, F. The functional SNPs picked out in the two cells in DiR-seq analysis were listed in **Table S5**. Notably, the functional SNPs discovered in the two cell lines displayed great diversity. On the other hand, the SNP rs10187424 exhibited gene regulatory activity in both two cell lines. It indicated that the functions and mechanisms of risk SNPs might be highly cell-specific.

**Figure 5.**
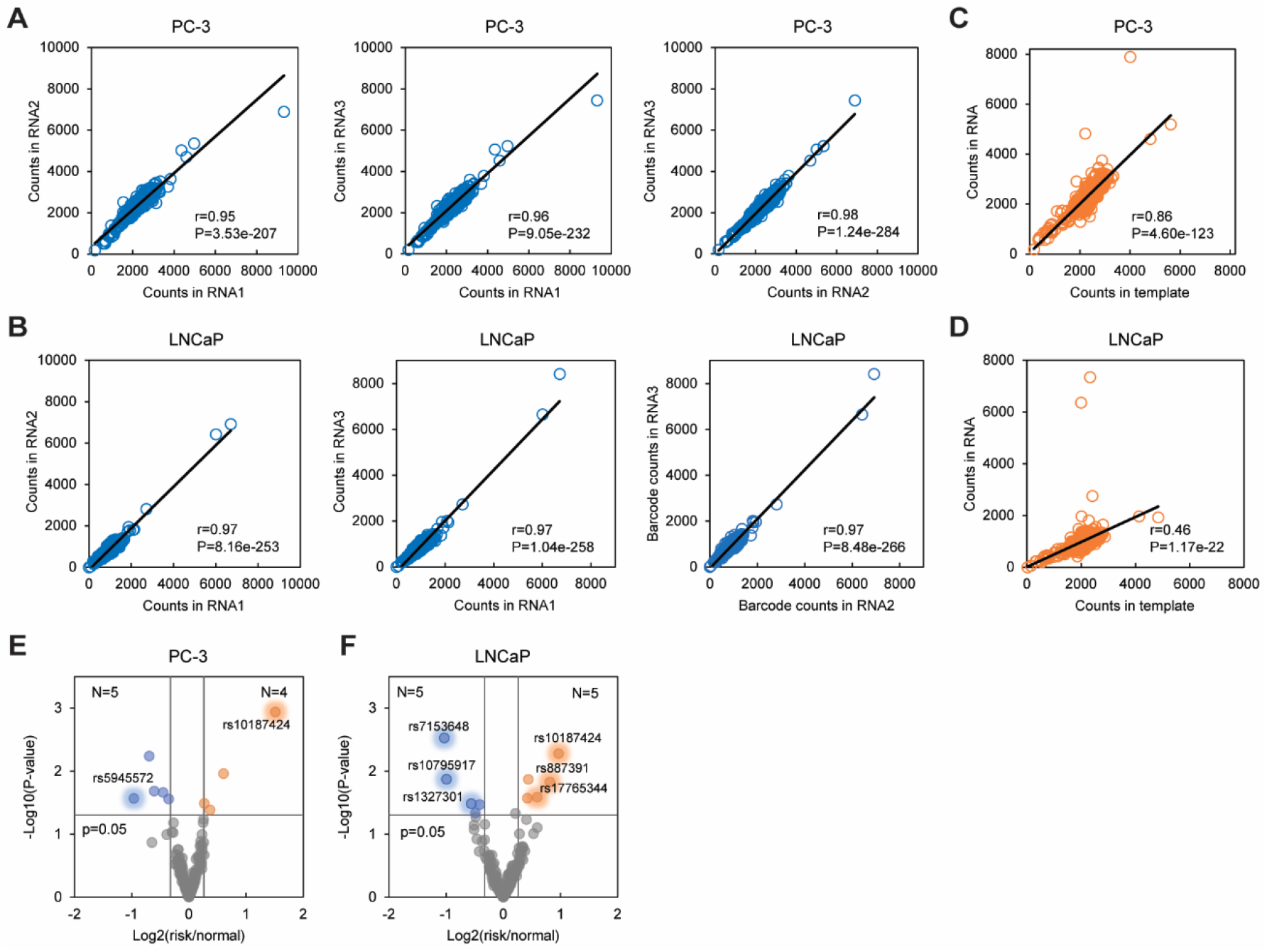
DiR-seq assays discovered regulatory SNPs in prostate cancer cells. (**A**) Consistency evaluation between individual biological replicates of DiR-seq assay of 213 prostate cancer risk SNPs in PC-3 cells. (**B**) Consistency evaluation between individual biological replicates of DiR-seq assay of 213 prostate cancer risk SNPs in LNCaP cells. (**C**) Scatter plot of DiR-seq tag counts in RNA and template in PC-3 cells. (**D**) Scatter plot of DiR-seq tag counts in RNA and template in LNCaP cells. The correlation coefficient values and P values were calculated with Pearson correlation analysis in A-D. (**E**) Volcano plot of DiR-seq analysis results in the PC-3 cell line of allele ratio.P values came from a two-tailed Student’s t-test of the expression of individual alleles. The blue dots represent SNP sites satisfying the criteria of fold change< 0.8, P<0.05, and the orange dots represent SNP sites satisfying the criteria of fold change>1.2, P<0.05. (**F**) Volcano plot of DiR-seq analysis results in LNCaP cell line of allele ratio. P values came from a two-tailed Student’s t-test of the expression of individual alleles. The blue dots represent SNP sites satisfying the criteria of fold change< 0.8, P<0.05, and the orange dots represent SNP sites satisfying the criteria of fold change>1.2, P<0.05.

### Transcription regulatory activity of selected SNPs

Specifically, for rs5945572, the risk ‘A’ allele showed significantly decreased transcriptional activity compared to the normal ‘G’ allele (**Fig. 6A**), and for rs10187424, the risk ‘T’ allele showed significantly increased transcriptional activity compared to the normal ‘C’ allele in PC-3 cells (**Fig. 6B**). Six most functional SNPs with a significant allelic difference were picked out in DiR-seq of LNCaP cells. For the three sites rs10795917, rs7153648, and rs1327301, risk alleles showed significantly lower transcriptional activity relative to the normal allele (**Fig. 6C, D, F**). And for the sites rs17765344, rs10187424, and rs887391, risk alleles showed significantly higher transcriptional activity relative to the normal allele (**Fig. 6E, G, H**). Notably, the SNP rs887391 exhibited similar allele-specific transcriptional activity in both PC-3 and LNCaP cells, for which the biological function and underlying mechanism had been illustrated previously[10,28]. It indicated that other functional SNPs we highlighted have potential research value in prostate cancer risk diseases.

**Figure 6.**
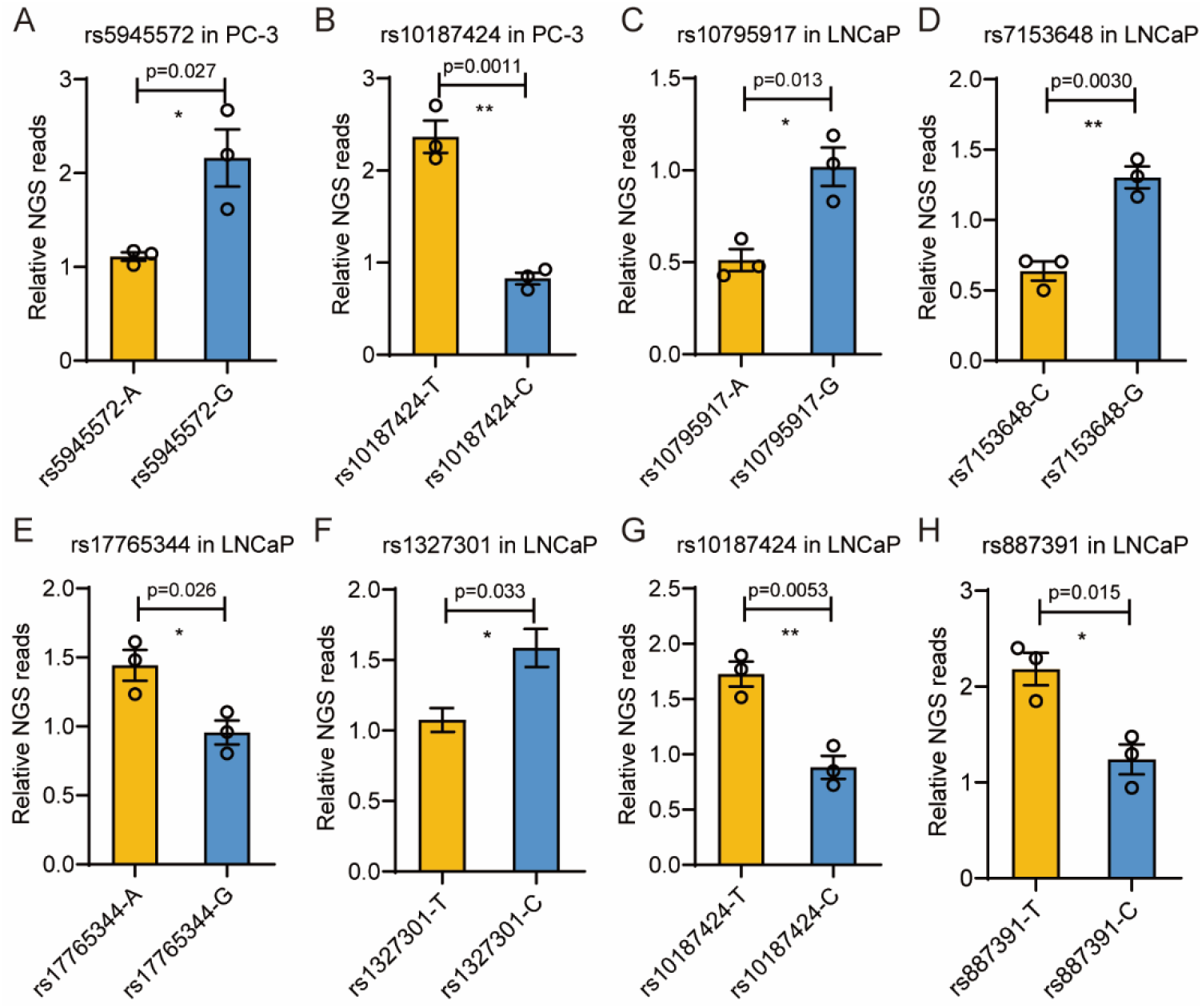
Reporter express levels of selected regulatory SNPs. (**A**) Reporter gene expression level for regulatory SNP rs5945572 alleles in the DiR-seq analysis in PC-3 cells. (**B**) Reporter gene expression level for regulatory SNP rs10187424 alleles in the DiR-seq analysis in PC-3 cells. (**C**) Reporter gene expression level for regulatory SNP rs10795917 alleles in the DiR-seq analysis in LNCaP cells. (**D**) Reporter gene expression level for regulatory SNP rs7153648 alleles in the DiR-seq analysis in LNCaP cells. (**E**) Reporter gene expression level for regulatory SNP rs17765344 alleles in the DiR-seq analysis in LNCaP cells. (**F**) Reporter gene expression level for regulatory SNP rs1327301 alleles in the DiR-seq analysis in LNCaP cells. (**G**) Reporter gene expression level for regulatory SNP rs10187424 alleles in the DiR-seq analysis in LNCaP cells. (**H**) Reporter gene expression level for regulatory SNP rs887391 alleles in the DiR-seq analysis in LNCaP cells. Risk alleles in orange and non-risk alleles in blue. Mean ± SEM of three independent experiments. *p<0.05, **p<0.01, two-tailed Student’s t-test.

## Discussion

Here, we developed a dinucleotide-tag reporter assay (DiR) system and applied it in screening the causal prostate cancer risk SNPs that possess potential gene regulatory functions. The DiR system has several advantages over the present reporter gene assay systems. Firstly, the DiR system employs dinucleotide barcodes to avoid nucleotide composition bias, which comes with the common 10-20nt barcodes and has to be minimized using multiple tags for each regulatory element [18–21]. The lower tag composition bias makes the DiR system more accurate and suitable to investigate the tiny difference between risk and normal alleles in regulating gene expression. Secondly, the DiR reporter system allows reporter assay through both Next-Generation sequencing and general throughput qPCR analysis, making the reporter assay system versatile. Thirdly, compared with the traditional luciferase reporter assay system, the DiR system holds a higher mRNA expression level, making it more stable and robust for moderate activity regulatory elements such as risk SNPs. Fourthly, like all the previous massive parallel methods, the DiR system performs reporter assay in one single transfection and does not need transfection normalization between tests like what happens in the luciferase activity assay. The normalization itself by *Renilla* luciferase activity can make the assay results complicated, especially for time-course investigations. Collectively, these advantages will hopefully make the DiR system suitable for broad application, especially for the functional study of regulatory SNPs.

Unavoidably, The DiR system also has limitations in some aspects. Firstly, the DiR system only has 628 dinucleotide tags, which is hardly high-throughput and limited for the massive parallel analysis. Secondly, even though barcoded using dinucleotide, the tag itself still might bring about minimal bias in cells not tested in this work. Finally, the DiR reporter plasmids have to be constructed individually for each regulatory element, making it labor-intensive and time-consuming like the luciferase reporter assay.

The DiR-seq assays for the prostate cancer risk SNPs in PC-3 and LNCaP cells identified gene regulatory SNP sites, and they exhibited high diversity in the two cell lines. Given that these two cell lines originate from different metastasis sites and different cancer patients and are largely different in many aspects, such as androgen dependency and gene expression profile[29,30], it indicates that the regulatory SNPs are supposed to function in a highly cell-specific manner. Therefore, to study the biological function and mechanism of risk SNPs, employing appropriate cancer cell lines is necessary. Overall, our DiR system provides a versatile reporter gene assay method for dissecting the gene regulatory effects of risk SNPs and can be easily applied to risk SNPs for other cancers.

In general, we developed a versatile DiR assay system that has multiple advantages over the traditional DNA barcode-based reporter system and the luciferase reporter assay. We identified regulatory prostate cancer risk SNPs by DiR-seq analysis in two prostate cancer cell lines and elucidated the allelic report activity. The results described here should be valuable for further studies on the biological function and underlying mechanism for the functional SNPs. Besides, wide applications of DiR assay in other cancer may identify more functional cancer risk SNPs.

## Supporting information

Supplementary Sequences

## Data and Code Availability

The raw sequence data generated using the Illumina Hiseq-PE150 platform for DiR-seq and 10-nucleotide tag reporter assay have been publicly available in the Gene Expression Omnibus (GEO) database under the accession number GSE165765.

## Acknowledgments

We thank Caiyun Sun, Xiangmei Ren, and Rui Wang from the State Key Laboratory of Microbial Technology of Shandong University for assistance with EnSpire multimode plate reader and M200 PRO multimode plate reader. This work was funded by the National Natural Science Foundation of China [grant numbers 31872809], Shandong Provincial Natural Science Foundation, China (ZR2016CM50), and Qilu Young Scholar to Q.H.

## CRediT authorship contribution statement

Naixia Ren: Data curation, Formal analysis, Investigation, Methodology, Visualization, Writing – original draft. Bo Li: Investigation, Methodology, Visualization, Writing – original draft. Qingqing Liu: Investigation, Methodology, Validation, Visualization, Writing – review & editing. Lele Yang: Methodology, Visualization, Writing – review & editing. Xiaodan Liu: Methodology, Visualization, Writing review & editing. Qilai Huang: Conceptualization, Project administration, Supervision, Writing – review & editing, Funding acquisition.

