## Supplementary Sequences for "A dinucleotide tag-based parallel reporter gene assay method"

**SUPPLEMENTAL SEQUENCES**

**DiR001:**

Tgcatctcaattagtcagcaaccatagtcccgcccctaactccgcccatcccgcccctaactccgcccagttccgcccattctccgccccatcgctgactaattttttttatttatgcagaggccgaggccgcctcggcctctgagctattccagaagtagtgaggaggcttttttggaggcctaggcttttgcaaaaagcttggcattccggtactgttggtaaagccacctgagtgaccgtgcgaggtcagtacagtcgttccagtgctcgaagcgaaggtcgtggatctgcctaccggtagaacgctgtgcgttgatcagagaggcgaactgcgtgtgagtggtcctatgactacgtccggttaggtacacagtcgcgaagcgaccaacgccttgatcgaccaggagtgatgcctgcactctgcagacctagcttgctgcgacgaagacgcacactccttcatcgttgaccgcctgaagtctctgactgagtcctgaggctaccaggtgactagtgctgagttggactccatcttgctccagcactagagcatcctcgactcaggtgtcgcaggtccttaggacgatgacgctggtgcacttcctgacgtcgttgtcgatctggagcacggatagacgatgacgtagtgagagatcgtggctgacgtcgccagtcgagtgacaactgcgactcagttgcgctgaggagttgtgcttgtggacgaagtaccgacaggtctgaccgagacactcgacgcaagacgagccagtgagatcctcatacaggccaagcagtgcggacagatcgtgctgtaa

**SV40 promoter**；**optimized 450bp sequence**

**DiR002:**

TgcatctcaattagtcagcaaccatagtcccgcccctaactccgcccatcccgcccctaactccgcccagttccgcccattctccgccccatcgctgactaattttttttatttatgcagaggccgaggccgcctcggcctctgagctattccagaagtagtgaggaggcttttttggaggcctaggcttttgcaaaaagcttggcattccggtactgttggtaaagccacctgagtgaccgtgcgaggtcagtacagtcgttccagtgGGcgaagcgaaggtcgtggatctgcctaccggtagaacgctgtgcgttgatcagagaggcgaactgcgtgtgagtggtcctatgactacgtccggttaggtacacagtcgcgaagcgaccaacgccttgatcgaccaggagtgatgcctgcactctgcagacctagcttgctgcgacgaagacgcacactccttcatcgttgaccgcctgaagtctctgactgagtcctgaggctaccaggtgactagtgctgagttggactccatcttgctccagcactagagcatcctcgactcaggtgtcgcaggtccttaggacgatgacgctggtgcacttcctgacgtcgttgtcgatctggagcacggatagacgatgacgtagtgagagatcgtggctgacgtcgccagtcgagtgacaactgcgactcagttgcgctgaggagttgtgcttgtggacgaagtaccgacaggtctgaccgagacactcgacgcaagacgagccagtgagatcctcatacaggccaagcagtgcggacagatcgtgctgtaa

**DiR003: GTACAGTCGTTCCAGTGTACGAAGCGAAGGTCGTG**

**DiR004: GTACAGTCGTTCCAGTGACCGAAGCGAAGGTCGTG**

**DiR005: TACAGTCGTTCCAGTGCTGAAAGCGAAGGTCGTGGATC**

**DiR006: TACAGTCGTTCCAGTGCTTCAAGCGAAGGTCGTGGATC**

**DiR007: TACAGTCGTTCCAGTGCTATAAGCGAAGGTCGTGGATC**

**DiR008: CAGTCGTTCCAGTGCTCGCCGCGAAGGTCGTGGATCTG**

**DiR009: CAGTCGTTCCAGTGCTCGGTGCGAAGGTCGTGGATCTG**

**DiR010: CAGTCGTTCCAGTGCTCGTGGCGAAGGTCGTGGATCTG**

**DiR011: GTCGTTCCAGTGCTCGAACTGAAGGTCGTGGATCTGCC**

**DiR012: GTCGTTCCAGTGCTCGAATAGAAGGTCGTGGATCTGCC**

**DiR013: GTCGTTCCAGTGCTCGAAAGGAAGGTCGTGGATCTGCC**

**DiR014: CGTTCCAGTGCTCGAAGCCCAGGTCGTGGATCTGCCTA**

**DiR015: CGTTCCAGTGCTCGAAGCTGAGGTCGTGGATCTGCCTA**

**DiR016: CGTTCCAGTGCTCGAAGCATAGGTCGTGGATCTGCCTA**

**DiR017: TTCCAGTGCTCGAAGCGACCGTCGTGGATCTGCCTACC**

**DiR018: TTCCAGTGCTCGAAGCGAGTGTCGTGGATCTGCCTACC**

**DiR019: TTCCAGTGCTCGAAGCGATAGTCGTGGATCTGCCTACC**

**DiR020: CCAGTGCTCGAAGCGAAGCCCGTGGATCTGCCTACCGG**

**DiR021: CCAGTGCTCGAAGCGAAGTGCGTGGATCTGCCTACCGG**

**DiR022: CCAGTGCTCGAAGCGAAGAACGTGGATCTGCCTACCGG**

**DiR023: AGTGCTCGAAGCGAAGGTGATGGATCTGCCTACCGGTA**

**DiR024: AGTGCTCGAAGCGAAGGTTCTGGATCTGCCTACCGGTA**

**DiR025: AGTGCTCGAAGCGAAGGTATTGGATCTGCCTACCGGTA**

**DiR026: TGCTCGAAGCGAAGGTCGCCGATCTGCCTACCGGTAGA**

**DiR027: TGCTCGAAGCGAAGGTCGGTGATCTGCCTACCGGTAGA**

**DiR028: TGCTCGAAGCGAAGGTCGAAGATCTGCCTACCGGTAGA**

**DiR029: CTCGAAGCGAAGGTCGTGCCTCTGCCTACCGGTAGAAC**

**DiR030: CTCGAAGCGAAGGTCGTGTGTCTGCCTACCGGTAGAAC**

**DiR031: CTCGAAGCGAAGGTCGTGATTCTGCCTACCGGTAGAAC**

**DiR032: CGAAGCGAAGGTCGTGGACATGCCTACCGGTAGAACGC**

**DiR033: CGAAGCGAAGGTCGTGGAGTTGCCTACCGGTAGAACGC**

**DiR034: CGAAGCGAAGGTCGTGGAAGTGCCTACCGGTAGAACGC**

**DiR035: AAGCGAAGGTCGTGGATCCCCCTACCGGTAGAACGCTG**

**DiR036: AAGCGAAGGTCGTGGATCGTCCTACCGGTAGAACGCTG**

**DiR037: AAGCGAAGGTCGTGGATCAACCTACCGGTAGAACGCTG**

**DiR038: GCGAAGGTCGTGGATCTGGTTACCGGTAGAACGCTGTG**

**DiR039: GCGAAGGTCGTGGATCTGTGTACCGGTAGAACGCTGTG**

**DiR040: GCGAAGGTCGTGGATCTGAATACCGGTAGAACGCTGTG**

**DiR041: GAAGGTCGTGGATCTGCCCCCCGGTAGAACGCTGTGCG**

**DiR042: GAAGGTCGTGGATCTGCCGTCCGGTAGAACGCTGTGCG**

**DiR043: GAAGGTCGTGGATCTGCCAGCCGGTAGAACGCTGTGCG**

**DiR044: AGGTCGTGGATCTGCCTAGTGGTAGAACGCTGTGCGTT**

**DiR045: AGGTCGTGGATCTGCCTATGGGTAGAACGCTGTGCGTT**

**DiR046: AGGTCGTGGATCTGCCTAAAGGTAGAACGCTGTGCGTT**

**DiR047: GTCGTGGATCTGCCTACCCCTAGAACGCTGTGCGTTGA**

**DiR048: GTCGTGGATCTGCCTACCTATAGAACGCTGTGCGTTGA**

**DiR049: GTCGTGGATCTGCCTACCATTAGAACGCTGTGCGTTGA**

**DiR050: CGTGGATCTGCCTACCGGCCGAACGCTGTGCGTTGATC**

**DiR051: CGTGGATCTGCCTACCGGGTGAACGCTGTGCGTTGATC**

**DiR052: CGTGGATCTGCCTACCGGAGGAACGCTGTGCGTTGATC**

**DiR053: TGGATCTGCCTACCGGTACCACGCTGTGCGTTGATCAG**

**DiR054: TGGATCTGCCTACCGGTATGACGCTGTGCGTTGATCAG**

**DiR055: TGGATCTGCCTACCGGTAATACGCTGTGCGTTGATCAG**

**DiR056: GGATCTGCCTACCGGTAGACTGCTGTGCGTTGATCAGAGA**

**DiR057: GGATCTGCCTACCGGTAGAGGGCTGTGCGTTGATCAGAGA**

**DiR058: GGATCTGCCTACCGGTAGATAGCTGTGCGTTGATCAGAGA**

**DiR059: TCTGCCTACCGGTAGAACCTTGTGCGTTGATCAGAGAG**

**DiR060: TCTGCCTACCGGTAGAACTATGTGCGTTGATCAGAGAG**

**DiR061: TCTGCCTACCGGTAGAACAGTGTGCGTTGATCAGAGAG**

**DiR062: TGCCTACCGGTAGAACGCCCTGCGTTGATCAGAGAGGC**

**DiR063: TGCCTACCGGTAGAACGCGTTGCGTTGATCAGAGAGGC**

**DiR064: TGCCTACCGGTAGAACGCAATGCGTTGATCAGAGAGGC**

**DiR065: CCTACCGGTAGAACGCTGCCCGTTGATCAGAGAGGCGA**

**DiR066: CCTACCGGTAGAACGCTGGTCGTTGATCAGAGAGGCGA**

**DiR067: CCTACCGGTAGAACGCTGAACGTTGATCAGAGAGGCGA**

**DiR068: TACCGGTAGAACGCTGTGGATTGATCAGAGAGGCGAAC**

**DiR069: TACCGGTAGAACGCTGTGTCTTGATCAGAGAGGCGAAC**

**DiR070: TACCGGTAGAACGCTGTGATTTGATCAGAGAGGCGAAC**

**DiR071: CCGGTAGAACGCTGTGCGCCGATCAGAGAGGCGAACTG**

**DiR072: CCGGTAGAACGCTGTGCGGGGATCAGAGAGGCGAACTG**

**DiR073: CCGGTAGAACGCTGTGCGAAGATCAGAGAGGCGAACTG**

**DiR074: GGTAGAACGCTGTGCGTTCCTCAGAGAGGCGAACTGCG**

**DiR075: GGTAGAACGCTGTGCGTTTGTCAGAGAGGCGAACTGCG**

**DiR076: GGTAGAACGCTGTGCGTTATTCAGAGAGGCGAACTGCG**

**DiR077: TAGAACGCTGTGCGTTGACAAGAGAGGCGAACTGCGTG**

**DiR078: TAGAACGCTGTGCGTTGAGTAGAGAGGCGAACTGCGTG**

**DiR079: TAGAACGCTGTGCGTTGAAGAGAGAGGCGAACTGCGTG**

**DiR080: GAACGCTGTGCGTTGATCCCAGAGGCGAACTGCGTGTG**

**DiR081: GAACGCTGTGCGTTGATCGTAGAGGCGAACTGCGTGTG**

**DiR082: GAACGCTGTGCGTTGATCTAAGAGGCGAACTGCGTGTG**

**DiR083: ACGCTGTGCGTTGATCAGCCAGGCGAACTGCGTGTGAG**

**DiR084: ACGCTGTGCGTTGATCAGGTAGGCGAACTGCGTGTGAG**

**DiR085: ACGCTGTGCGTTGATCAGTAAGGCGAACTGCGTGTGAG**

**DiR086: GCTGTGCGTTGATCAGAGCCGCGAACTGCGTGTGAGTG**

**DiR087: GCTGTGCGTTGATCAGAGGTGCGAACTGCGTGTGAGTG**

**DiR088: GCTGTGCGTTGATCAGAGTAGCGAACTGCGTGTGAGTG**

**DiR089: TGTGCGTTGATCAGAGAGCTGAACTGCGTGTGAGTGGT**

**DiR090: TGTGCGTTGATCAGAGAGTAGAACTGCGTGTGAGTGGT**

**DiR091: TGTGCGTTGATCAGAGAGAGGAACTGCGTGTGAGTGGT**

**DiR092: TGCGTTGATCAGAGAGGCCCACTGCGTGTGAGTGGTCC**

**DiR093: TGCGTTGATCAGAGAGGCTGACTGCGTGTGAGTGGTCC**

**DiR094: TGCGTTGATCAGAGAGGCATACTGCGTGTGAGTGGTCC**

**DiR095: CGTTGATCAGAGAGGCGACTTGCGTGTGAGTGGTCCTA**

**DiR096: CGTTGATCAGAGAGGCGAGGTGCGTGTGAGTGGTCCTA**

**DiR097: CGTTGATCAGAGAGGCGATATGCGTGTGAGTGGTCCTA**

**DiR098: TTGATCAGAGAGGCGAACCCCGTGTGAGTGGTCCTATG**

**DiR099: TTGATCAGAGAGGCGAACGTCGTGTGAGTGGTCCTATG**

**DiR100: TTGATCAGAGAGGCGAACAACGTGTGAGTGGTCCTATG**

**DiR101: GATCAGAGAGGCGAACTGGATGTGAGTGGTCCTATGAC**

**DiR102: GATCAGAGAGGCGAACTGTCTGTGAGTGGTCCTATGAC**

**DiR103: GATCAGAGAGGCGAACTGATTGTGAGTGGTCCTATGAC**

**DiR104: TCAGAGAGGCGAACTGCGCCTGAGTGGTCCTATGACTA**

**DiR105: TCAGAGAGGCGAACTGCGGTTGAGTGGTCCTATGACTA**

**DiR106: TCAGAGAGGCGAACTGCGAATGAGTGGTCCTATGACTA**

**DiR107: AGAGAGGCGAACTGCGTGCCAGTGGTCCTATGACTACG**

**DiR108: AGAGAGGCGAACTGCGTGGTAGTGGTCCTATGACTACG**

**DiR109: AGAGAGGCGAACTGCGTGAAAGTGGTCCTATGACTACG**

**DiR110: AGAGGCGAACTGCGTGTGCCTGGTCCTATGACTACGTC**

**DiR111: AGAGGCGAACTGCGTGTGGTTGGTCCTATGACTACGTC**

**DiR112: AGAGGCGAACTGCGTGTGTATGGTCCTATGACTACGTC**

**DiR113: AGGCGAACTGCGTGTGAGCCGTCCTATGACTACGTCCG**

**DiR114: AGGCGAACTGCGTGTGAGGTGTCCTATGACTACGTCCG**

**DiR115: AGGCGAACTGCGTGTGAGAAGTCCTATGACTACGTCCG**

**DiR116: GCGAACTGCGTGTGAGTGCCCCTATGACTACGTCCGGT**

**DiR117: GCGAACTGCGTGTGAGTGTGCCTATGACTACGTCCGGT**

**DiR118: GCGAACTGCGTGTGAGTGAACCTATGACTACGTCCGGT**

**DiR119: GAACTGCGTGTGAGTGGTGTTATGACTACGTCCGGTTA**

**DiR120: GAACTGCGTGTGAGTGGTTGTATGACTACGTCCGGTTA**

**DiR121: GAACTGCGTGTGAGTGGTAATATGACTACGTCCGGTTA**

**DiR122: ACTGCGTGTGAGTGGTCCCCTGACTACGTCCGGTTAGG**

**DiR123: ACTGCGTGTGAGTGGTCCGTTGACTACGTCCGGTTAGG**

**DiR124: ACTGCGTGTGAGTGGTCCAGTGACTACGTCCGGTTAGG**

**DiR125: TGCGTGTGAGTGGTCCTACCACTACGTCCGGTTAGGTA**

**DiR126: TGCGTGTGAGTGGTCCTAGTACTACGTCCGGTTAGGTA**

**DiR127: TGCGTGTGAGTGGTCCTAAAACTACGTCCGGTTAGGTA**

**DiR128: CGTGTGAGTGGTCCTATGCTTACGTCCGGTTAGGTACA**

**DiR129: CGTGTGAGTGGTCCTATGGGTACGTCCGGTTAGGTACA**

**DiR130: CGTGTGAGTGGTCCTATGTATACGTCCGGTTAGGTACA**

**DiR131: TGTGAGTGGTCCTATGACCCCGTCCGGTTAGGTACACA**

**DiR132: TGTGAGTGGTCCTATGACGTCGTCCGGTTAGGTACACA**

**DiR133: TGTGAGTGGTCCTATGACAGCGTCCGGTTAGGTACACA**

**DiR134: TGAGTGGTCCTATGACTAGATCCGGTTAGGTACACAGT**

**DiR135: TGAGTGGTCCTATGACTATCTCCGGTTAGGTACACAGT**

**DiR136: TGAGTGGTCCTATGACTAATTCCGGTTAGGTACACAGT**

**DiR137: AGTGGTCCTATGACTACGCACGGTTAGGTACACAGTCG**

**DiR138: AGTGGTCCTATGACTACGGTCGGTTAGGTACACAGTCG**

**DiR139: AGTGGTCCTATGACTACGAGCGGTTAGGTACACAGTCG**

**DiR140: TGGTCCTATGACTACGTCGAGTTAGGTACACAGTCGCG**

**DiR141: TGGTCCTATGACTACGTCTCGTTAGGTACACAGTCGCG**

**DiR142: TGGTCCTATGACTACGTCATGTTAGGTACACAGTCGCG**

**DiR143: GTCCTATGACTACGTCCGCCTAGGTACACAGTCGCGAA**

**DiR144: GTCCTATGACTACGTCCGTGTAGGTACACAGTCGCGAA**

**DiR145: GTCCTATGACTACGTCCGAATAGGTACACAGTCGCGAA**

**DiR146: CCTATGACTACGTCCGGTCCGGTACACAGTCGCGAAGC**

**DiR147: CCTATGACTACGTCCGGTGTGGTACACAGTCGCGAAGC**

**DiR148: CCTATGACTACGTCCGGTAGGGTACACAGTCGCGAAGC**

**DiR149: TATGACTACGTCCGGTTACCTACACAGTCGCGAAGCGA**

**DiR150: TATGACTACGTCCGGTTATATACACAGTCGCGAAGCGA**

**DiR151: TATGACTACGTCCGGTTAATTACACAGTCGCGAAGCGA**

**DiR152: TGACTACGTCCGGTTAGGCCCACAGTCGCGAAGCGACC**

**DiR153: TGACTACGTCCGGTTAGGGTCACAGTCGCGAAGCGACC**

**DiR154: TGACTACGTCCGGTTAGGAGCACAGTCGCGAAGCGACC**

**DiR155: ACTACGTCCGGTTAGGTAGTCAGTCGCGAAGCGACCAA**

**DiR156: ACTACGTCCGGTTAGGTATCCAGTCGCGAAGCGACCAA**

**DiR157: ACTACGTCCGGTTAGGTAAGCAGTCGCGAAGCGACCAA**

**DiR158: TACGTCCGGTTAGGTACAGTGTCGCGAAGCGACCAACG**

**DiR159: TACGTCCGGTTAGGTACATCGTCGCGAAGCGACCAACG**

**DiR160: TACGTCCGGTTAGGTACAAGGTCGCGAAGCGACCAACG**

**DiR161: GTCCGGTTAGGTACACAAGCGCGAAGCGACCAACGC**

**DiR162: GTCCGGTTAGGTACACACCCGCGAAGCGACCAACGC**

**DiR163: GTCCGGTTAGGTACACATACGCGAAGCGACCAACGC**

**DiR164: CCGGTTAGGTACACAGTTCCGAAGCGACCAACGCCT**

**DiR165: CCGGTTAGGTACACAGTGACGAAGCGACCAACGCCT**

**DiR166: CCGGTTAGGTACACAGTATCGAAGCGACCAACGCCT**

**DiR167: GGTTAGGTACACAGTCGTCAAGCGACCAACGCCTTG**

**DiR168: GGTTAGGTACACAGTCGGAAAGCGACCAACGCCTTG**

**DiR169: GGTTAGGTACACAGTCGATAAGCGACCAACGCCTTG**

**DiR170: TTAGGTACACAGTCGCGGGGCGACCAACGCCTTGAT**

**DiR171: TTAGGTACACAGTCGCGCCGCGACCAACGCCTTGAT**

**DiR172: TTAGGTACACAGTCGCGTTGCGACCAACGCCTTGAT**

**DiR173: AGGTACACAGTCGCGAAAGGACCAACGCCTTGATCG**

**DiR174: AGGTACACAGTCGCGAATAGACCAACGCCTTGATCG**

**DiR175: AGGTACACAGTCGCGAACTGACCAACGCCTTGATCG**

**DiR176: GTACACAGTCGCGAAGCTGCCAACGCCTTGATCGAC**

**DiR177: GTACACAGTCGCGAAGCACCCAACGCCTTGATCGAC**

**DiR178: GTACACAGTCGCGAAGCCTCCAACGCCTTGATCGAC**

**DiR179: ACACAGTCGCGAAGCGAGGAACGCCTTGATCGACCA**

**DiR180: ACACAGTCGCGAAGCGATAAACGCCTTGATCGACCA**

**DiR181: ACACAGTCGCGAAGCGAATAACGCCTTGATCGACCA**

**DiR182: ACAGTCGCGAAGCGACCGGCGCCTTGATCGACCAGG**

**DiR183: ACAGTCGCGAAGCGACCCCCGCCTTGATCGACCAGG**

**DiR184: ACAGTCGCGAAGCGACCTTCGCCTTGATCGACCAGG**

**DiR185: AGTCGCGAAGCGACCAATCCCTTGATCGACCAGGAG**

**DiR186: AGTCGCGAAGCGACCAAGACCTTGATCGACCAGGAG**

**DiR187: AGTCGCGAAGCGACCAAATCCTTGATCGACCAGGAG**

**DiR188: TCGCGAAGCGACCAACGGGTTGATCGACCAGGAGTG**

**DiR189: TCGCGAAGCGACCAACGTATTGATCGACCAGGAGTG**

**DiR190: TCGCGAAGCGACCAACGATTTGATCGACCAGGAGTG**

**DiR191: GCGAAGCGACCAACGCCGGGATCGACCAGGAGTGAT**

**DiR192: GCGAAGCGACCAACGCCACGATCGACCAGGAGTGAT**

**DiR193: GCGAAGCGACCAACGCCCAGATCGACCAGGAGTGAT**

**DiR194: GAAGCGACCAACGCCTTTGTCGACCAGGAGTGATGC**

**DiR195: GAAGCGACCAACGCCTTACTCGACCAGGAGTGATGC**

**DiR196: GAAGCGACCAACGCCTTCTTCGACCAGGAGTGATGC**

**DiR197: AGCGACCAACGCCTTGAGGGACCAGGAGTGATGCCT**

**DiR198: AGCGACCAACGCCTTGACAGACCAGGAGTGATGCCT**

**DiR199: AGCGACCAACGCCTTGAATGACCAGGAGTGATGCCT**

**DiR200: CGACCAACGCCTTGATCTGCCAGGAGTGATGCCTGC**

**DiR201: CGACCAACGCCTTGATCACCCAGGAGTGATGCCTGC**

**DiR202: CGACCAACGCCTTGATCCTCCAGGAGTGATGCCTGC**

**DiR203: ACCAACGCCTTGATCGAGGAGGAGTGATGCCTGCAC**

**DiR204: ACCAACGCCTTGATCGATAAGGAGTGATGCCTGCAC**

**DiR205: ACCAACGCCTTGATCGAATAGGAGTGATGCCTGCAC**

**DiR206: CAACGCCTTGATCGACCCCGAGTGATGCCTGCACTC**

**DiR207: CAACGCCTTGATCGACCTAGAGTGATGCCTGCACTC**

**DiR208: CAACGCCTTGATCGACCGTGAGTGATGCCTGCACTC**

**DiR209: ACGCCTTGATCGACCAGTGGTGATGCCTGCACTCTG**

**DiR210: ACGCCTTGATCGACCAGACGTGATGCCTGCACTCTG**

**DiR211: ACGCCTTGATCGACCAGCTGTGATGCCTGCACTCTG**

**DiR212: GCCTTGATCGACCAGGAAGGATGCCTGCACTCTGCA**

**DiR213: GCCTTGATCGACCAGGACCGATGCCTGCACTCTGCA**

**DiR214: GCCTTGATCGACCAGGATAGATGCCTGCACTCTGCA**

**DiR215: CTTGATCGACCAGGAGTTGTGCCTGCACTCTGCAGA**

**DiR216: CTTGATCGACCAGGAGTACTGCCTGCACTCTGCAGA**

**DiR217: CTTGATCGACCAGGAGTCTTGCCTGCACTCTGCAGA**

**DiR218: TGATCGACCAGGAGTGAACCCTGCACTCTGCAGACC**

**DiR219: TGATCGACCAGGAGTGAGACCTGCACTCTGCAGACC**

**DiR220: TGATCGACCAGGAGTGACTCCTGCACTCTGCAGACC**

**DiR221: ATCGACCAGGAGTGATGGGTGCACTCTGCAGACCTA**

**DiR222: ATCGACCAGGAGTGATGTATGCACTCTGCAGACCTA**

**DiR223: ATCGACCAGGAGTGATGATTGCACTCTGCAGACCTA**

**DiR224: CGACCAGGAGTGATGCCACCACTCTGCAGACCTAGC**

**DiR225: CGACCAGGAGTGATGCCGACACTCTGCAGACCTAGC**

**DiR226: CGACCAGGAGTGATGCCCTCACTCTGCAGACCTAGC**

**DiR227: ACCAGGAGTGATGCCTGGGCTCTGCAGACCTAGCTT**

**DiR228: ACCAGGAGTGATGCCTGACCTCTGCAGACCTAGCTT**

**DiR229: ACCAGGAGTGATGCCTGTTCTCTGCAGACCTAGCTT**

**DiR230: CAGGAGTGATGCCTGCAGGCTGCAGACCTAGCTTGC**

**DiR231: CAGGAGTGATGCCTGCAACCTGCAGACCTAGCTTGC**

**DiR232: CAGGAGTGATGCCTGCATACTGCAGACCTAGCTTGC**

**DiR233: GGAGTGATGCCTGCACTGGGCAGACCTAGCTTGCTG**

**DiR234: GGAGTGATGCCTGCACTACGCAGACCTAGCTTGCTG**

**DiR235: GGAGTGATGCCTGCACTTAGCAGACCTAGCTTGCTG**

**DiR236: AGTGATGCCTGCACTCTAGAGACCTAGCTTGCTGCG**

**DiR237: AGTGATGCCTGCACTCTTAAGACCTAGCTTGCTGCG**

**DiR238: AGTGATGCCTGCACTCTCTAGACCTAGCTTGCTGCG**

**DiR239: TGATGCCTGCACTCTGCCCACCTAGCTTGCTGCGAC**

**DiR240: TGATGCCTGCACTCTGCTAACCTAGCTTGCTGCGAC**

**DiR241: TGATGCCTGCACTCTGCGTACCTAGCTTGCTGCGAC**

**DiR242: ATGCCTGCACTCTGCAGGGCTAGCTTGCTGCGACGA**

**DiR243: ATGCCTGCACTCTGCAGCACTAGCTTGCTGCGACGA**

**DiR244: ATGCCTGCACTCTGCAGTTCTAGCTTGCTGCGACGA**

**DiR245: GCCTGCACTCTGCAGACGGAGCTTGCTGCGACGAAG**

**DiR246: GCCTGCACTCTGCAGACACAGCTTGCTGCGACGAAG**

**DiR247: GCCTGCACTCTGCAGACTAAGCTTGCTGCGACGAAG**

**DiR248: CTGCACTCTGCAGACCTCCCTTGCTGCGACGAAGAC**

**DiR249: CTGCACTCTGCAGACCTTACTTGCTGCGACGAAGAC**

**DiR250: CTGCACTCTGCAGACCTGTCTTGCTGCGACGAAGAC**

**DiR251: GCACTCTGCAGACCTAGGGTGCTGCGACGAAGACGC**

**DiR252: GCACTCTGCAGACCTAGACTGCTGCGACGAAGACGC**

**DiR253: GCACTCTGCAGACCTAGTATGCTGCGACGAAGACGC**

**DiR254: ACTCTGCAGACCTAGCTACCTGCGACGAAGACGCAC**

**DiR255: ACTCTGCAGACCTAGCTGACTGCGACGAAGACGCAC**

**DiR256: ACTCTGCAGACCTAGCTCTCTGCGACGAAGACGCAC**

**DiR257: TCTGCAGACCTAGCTTGGGGCGACGAAGACGCACAC**

**DiR258: TCTGCAGACCTAGCTTGACGCGACGAAGACGCACAC**

**DiR259: TCTGCAGACCTAGCTTGTAGCGACGAAGACGCACAC**

**DiR260: TGCAGACCTAGCTTGCTAGGACGAAGACGCACACTC**

**DiR261: TGCAGACCTAGCTTGCTTAGACGAAGACGCACACTC**

**DiR262: TGCAGACCTAGCTTGCTCTGACGAAGACGCACACTC**

**DiR263: CAGACCTAGCTTGCTGCTGCGAAGACGCACACTCCT**

**DiR264: CAGACCTAGCTTGCTGCACCGAAGACGCACACTCCT**

**DiR265: CAGACCTAGCTTGCTGCCTCGAAGACGCACACTCCT**

**DiR266: GACCTAGCTTGCTGCGATCAAGACGCACACTCCTTC**

**DiR267: GACCTAGCTTGCTGCGAGAAAGACGCACACTCCTTC**

**DiR268: GACCTAGCTTGCTGCGAATAAGACGCACACTCCTTC**

**DiR269: CCTAGCTTGCTGCGACGGGGACGCACACTCCTTCAT**

**DiR270: CCTAGCTTGCTGCGACGCCGACGCACACTCCTTCAT**

**DiR271: CCTAGCTTGCTGCGACGTTGACGCACACTCCTTCAT**

**DiR272: CTAGCTTGCTGCGACGAATGCGCACACTCCTTCATCGT**

**DiR273: CTAGCTTGCTGCGACGAAACCGCACACTCCTTCATCGT**

**DiR274: CTAGCTTGCTGCGACGAACTCGCACACTCCTTCATCGT**

**DiR275: GCTTGCTGCGACGAAGATCCACACTCCTTCATCGTT**

**DiR276: GCTTGCTGCGACGAAGAGACACACTCCTTCATCGTT**

**DiR277: GCTTGCTGCGACGAAGAATCACACTCCTTCATCGTT**

**DiR278: CTTGCTGCGACGAAGACGGGCACTCCTTCATCGTTGAC**

**DiR279: CTTGCTGCGACGAAGACGACCACTCCTTCATCGTTGAC**

**DiR280: CTTGCTGCGACGAAGACGTTCACTCCTTCATCGTTGAC**

**DiR281: GCTGCGACGAAGACGCAGGCTCCTTCATCGTTGACC**

**DiR282: GCTGCGACGAAGACGCAACCTCCTTCATCGTTGACC**

**DiR283: GCTGCGACGAAGACGCATTCTCCTTCATCGTTGACC**

**DiR284: TGCGACGAAGACGCACAGGCCTTCATCGTTGACCGC**

**DiR285: TGCGACGAAGACGCACAACCCTTCATCGTTGACCGC**

**DiR286: TGCGACGAAGACGCACATACCTTCATCGTTGACCGC**

**DiR287: CGACGAAGACGCACACTGGTTCATCGTTGACCGCCT**

**DiR288: CGACGAAGACGCACACTTATTCATCGTTGACCGCCT**

**DiR289: CGACGAAGACGCACACTATTTCATCGTTGACCGCCT**

**DiR290: ACGAAGACGCACACTCCGGCATCGTTGACCGCCTGA**

**DiR291: ACGAAGACGCACACTCCACCATCGTTGACCGCCTGA**

**DiR292: ACGAAGACGCACACTCCCACATCGTTGACCGCCTGA**

**DiR293: GAAGACGCACACTCCTTGGTCGTTGACCGCCTGAAG**

**DiR294: GAAGACGCACACTCCTTACTCGTTGACCGCCTGAAG**

**DiR295: GAAGACGCACACTCCTTTTTCGTTGACCGCCTGAAG**

**DiR296: AGACGCACACTCCTTCAGGGTTGACCGCCTGAAGTC**

**DiR297: AGACGCACACTCCTTCACAGTTGACCGCCTGAAGTC**

**DiR298: AGACGCACACTCCTTCAATGTTGACCGCCTGAAGTC**

**DiR299: ACGCACACTCCTTCATCAGTGACCGCCTGAAGTCTC**

**DiR300: ACGCACACTCCTTCATCCCTGACCGCCTGAAGTCTC**

**DiR301: ACGCACACTCCTTCATCTATGACCGCCTGAAGTCTC**

**DiR302: GCACACTCCTTCATCGTACACCGCCTGAAGTCTCTG**

**DiR303: GCACACTCCTTCATCGTGAACCGCCTGAAGTCTCTG**

**DiR304: GCACACTCCTTCATCGTCTACCGCCTGAAGTCTCTG**

**DiR305: ACACTCCTTCATCGTTGGGCGCCTGAAGTCTCTGAC**

**DiR306: ACACTCCTTCATCGTTGCACGCCTGAAGTCTCTGAC**

**DiR307: ACACTCCTTCATCGTTGTTCGCCTGAAGTCTCTGAC**

**DiR308: ACTCCTTCATCGTTGACTCCCTGAAGTCTCTGACTG**

**DiR309: ACTCCTTCATCGTTGACGACCTGAAGTCTCTGACTG**

**DiR310: CACTCCTTCATCGTTGACATCCTGAAGTCTCTGACTGA**

**DiR311: TCCTTCATCGTTGACCGGGTGAAGTCTCTGACTGAG**

**DiR312: CTCCTTCATCGTTGACCGTATGAAGTCTCTGACTGAGT**

**DiR313: CTCCTTCATCGTTGACCGATTGAAGTCTCTGACTGAGT**

**DiR314:** **CTACCAGGTGACTAGTGCCCAGTTGGACTCCATCTTGC**

**DiR315:** **CTACCAGGTGACTAGTGCGTAGTTGGACTCCATCTTGC**

**DiR316: CTACCAGGTGACTAGTGCAAAGTTGGACTCCATCTTGC**

**DiR317:** **ACCAGGTGACTAGTGCTGCCTTGGACTCCATCTTGCTC**

**DiR318:** **ACCAGGTGACTAGTGCTGGTTTGGACTCCATCTTGCTC**

**DiR319: ACCAGGTGACTAGTGCTGTATTGGACTCCATCTTGCTC**

**DiR320:** **CAGGTGACTAGTGCTGAGCCGGACTCCATCTTGCTCCA**

**DiR321:** **CAGGTGACTAGTGCTGAGGGGGACTCCATCTTGCTCCA**

**DiR322:** **CAGGTGACTAGTGCTGAGAAGGACTCCATCTTGCTCCA**

**DiR323: GGTGACTAGTGCTGAGTTCCACTCCATCTTGCTCCAGC**

**DiR324: GGTGACTAGTGCTGAGTTTAACTCCATCTTGCTCCAGC**

**DiR325:** **GGTGACTAGTGCTGAGTTATACTCCATCTTGCTCCAGC**

**DiR326: TGACTAGTGCTGAGTTGGCTTCCATCTTGCTCCAGCAC**

**DiR327:** **TGACTAGTGCTGAGTTGGGGTCCATCTTGCTCCAGCAC**

**DiR328:** **TGACTAGTGCTGAGTTGGTATCCATCTTGCTCCAGCAC**

**DiR329:** **ACTAGTGCTGAGTTGGACCACATCTTGCTCCAGCACTA**

**DiR330: ACTAGTGCTGAGTTGGACGTCATCTTGCTCCAGCACTA**

**DiR331: ACTAGTGCTGAGTTGGACAGCATCTTGCTCCAGCACTA**

**DiR332: TAGTGCTGAGTTGGACTCGTTCTTGCTCCAGCACTAGA**

**DiR333: TAGTGCTGAGTTGGACTCTCTCTTGCTCCAGCACTAGA**

**DiR334: TAGTGCTGAGTTGGACTCAGTCTTGCTCCAGCACTAGA**

**DiR335: GTGCTGAGTTGGACTCCACATTGCTCCAGCACTAGAGC**

**DiR336: GTGCTGAGTTGGACTCCAGTTTGCTCCAGCACTAGAGC**

**DiR337: GTGCTGAGTTGGACTCCAAGTTGCTCCAGCACTAGAGC**

**DiR338: GCTGAGTTGGACTCCATCCCGCTCCAGCACTAGAGCAT**

**DiR339: GCTGAGTTGGACTCCATCGGGCTCCAGCACTAGAGCAT**

**DiR340: GCTGAGTTGGACTCCATCAAGCTCCAGCACTAGAGCAT**

**DiR341: TGAGTTGGACTCCATCTTCTTCCAGCACTAGAGCATCC**

**DiR342: TGAGTTGGACTCCATCTTTATCCAGCACTAGAGCATCC**

**DiR343:** **TGAGTTGGACTCCATCTTAGTCCAGCACTAGAGCATCC**

**DiR344: AGTTGGACTCCATCTTGCCACAGCACTAGAGCATCCTC**

**DiR345: AGTTGGACTCCATCTTGCGTCAGCACTAGAGCATCCTC**

**DiR346: AGTTGGACTCCATCTTGCAGCAGCACTAGAGCATCCTC**

**DiR347: TTGGACTCCATCTTGCTCGTGCACTAGAGCATCCTCGA**

**DiR348: TTGGACTCCATCTTGCTCTCGCACTAGAGCATCCTCGA**

**DiR349: TTGGACTCCATCTTGCTCAGGCACTAGAGCATCCTCGA**

**DiR350: GGACTCCATCTTGCTCCACTACTAGAGCATCCTCGACT**

**DiR351: GGACTCCATCTTGCTCCATAACTAGAGCATCCTCGACT**

**DiR352: GGACTCCATCTTGCTCCAAGACTAGAGCATCCTCGACT**

**DiR353: ACTCCATCTTGCTCCAGCCTTAGAGCATCCTCGACTCA**

**DiR354: ACTCCATCTTGCTCCAGCGGTAGAGCATCCTCGACTCA**

**DiR355: ACTCCATCTTGCTCCAGCTATAGAGCATCCTCGACTCA**

**DiR356: TCCATCTTGCTCCAGCACCCGAGCATCCTCGACTCAGG**

**DiR357: TCCATCTTGCTCCAGCACGTGAGCATCCTCGACTCAGG**

**DiR358: TCCATCTTGCTCCAGCACAGGAGCATCCTCGACTCAGG**

**DiR359: CATCTTGCTCCAGCACTACCGCATCCTCGACTCAGGTG**

**DiR360: CATCTTGCTCCAGCACTATGGCATCCTCGACTCAGGTG**

**DiR361: CATCTTGCTCCAGCACTAATGCATCCTCGACTCAGGTG**

**DiR362: TCTTGCTCCAGCACTAGACTATCCTCGACTCAGGTGTC**

**DiR363: TCTTGCTCCAGCACTAGATAATCCTCGACTCAGGTGTC**

**DiR364: TCTTGCTCCAGCACTAGAAGATCCTCGACTCAGGTGTC**

**DiR365: TTGCTCCAGCACTAGAGCCCCCTCGACTCAGGTGTCGC**

**DiR366: TTGCTCCAGCACTAGAGCGACCTCGACTCAGGTGTCGC**

**DiR367: TTGCTCCAGCACTAGAGCTGCCTCGACTCAGGTGTCGC**

**DiR368: GCTCCAGCACTAGAGCATGTTCGACTCAGGTGTCGCAG**

**DiR369: GCTCCAGCACTAGAGCATTGTCGACTCAGGTGTCGCAG**

**DiR370: GCTCCAGCACTAGAGCATAATCGACTCAGGTGTCGCAG**

**DiR371: TCCAGCACTAGAGCATCCCAGACTCAGGTGTCGCAGGT**

**DiR372: TCCAGCACTAGAGCATCCGTGACTCAGGTGTCGCAGGT**

**DiR373: TCCAGCACTAGAGCATCCAGGACTCAGGTGTCGCAGGT**

**DiR374: CAGCACTAGAGCATCCTCCCCTCAGGTGTCGCAGGTCC**

**DiR375: CAGCACTAGAGCATCCTCTGCTCAGGTGTCGCAGGTCC**

**DiR376: CAGCACTAGAGCATCCTCATCTCAGGTGTCGCAGGTCC**

**DiR377: GCACTAGAGCATCCTCGAGGCAGGTGTCGCAGGTCCTT**

**DiR378: GCACTAGAGCATCCTCGATACAGGTGTCGCAGGTCCTT**

**DiR379: GCACTAGAGCATCCTCGAACCAGGTGTCGCAGGTCCTT**

**DiR380: ACTAGAGCATCCTCGACTGTGGTGTCGCAGGTCCTTAG**

**DiR381: ACTAGAGCATCCTCGACTTCGGTGTCGCAGGTCCTTAG**

**DiR382: ACTAGAGCATCCTCGACTAGGGTGTCGCAGGTCCTTAG**

**DiR383: TAGAGCATCCTCGACTCACCTGTCGCAGGTCCTTAGGA**

**DiR384: TAGAGCATCCTCGACTCATATGTCGCAGGTCCTTAGGA**

**DiR385: TAGAGCATCCTCGACTCAATTGTCGCAGGTCCTTAGGA**

**DiR386: GAGCATCCTCGACTCAGGCCTCGCAGGTCCTTAGGACG**

**DiR387: GAGCATCCTCGACTCAGGGTTCGCAGGTCCTTAGGACG**

**DiR388: GAGCATCCTCGACTCAGGAATCGCAGGTCCTTAGGACG**

**DiR389: GCATCCTCGACTCAGGTGCAGCAGGTCCTTAGGACGAT**

**DiR390: GCATCCTCGACTCAGGTGGTGCAGGTCCTTAGGACGAT**

**DiR391: GCATCCTCGACTCAGGTGAGGCAGGTCCTTAGGACGAT**

**DiR392: ATCCTCGACTCAGGTGTCCTAGGTCCTTAGGACGATGA**

**DiR393: ATCCTCGACTCAGGTGTCTAAGGTCCTTAGGACGATGA**

**DiR394: ATCCTCGACTCAGGTGTCAGAGGTCCTTAGGACGATGA**

**DiR395: CCTCGACTCAGGTGTCGCCCGTCCTTAGGACGATGACG**

**DiR396: CCTCGACTCAGGTGTCGCGTGTCCTTAGGACGATGACG**

**DiR397: CCTCGACTCAGGTGTCGCTAGTCCTTAGGACGATGACG**

**DiR398: TCGACTCAGGTGTCGCAGCCCCTTAGGACGATGACGCT**

**DiR399: TCGACTCAGGTGTCGCAGTGCCTTAGGACGATGACGCT**

**DiR400: TCGACTCAGGTGTCGCAGAACCTTAGGACGATGACGCT**

**DiR401: ACTCAGGTGTCGCAGGTGTTTAGGACGATGACGCTG**

**DiR402: ACTCAGGTGTCGCAGGTTGTTAGGACGATGACGCTG**

**DiR403: ACTCAGGTGTCGCAGGTAATTAGGACGATGACGCTG**

**DiR404: TCAGGTGTCGCAGGTCCCCAGGACGATGACGCTGGT**

**DiR405: TCAGGTGTCGCAGGTCCGGAGGACGATGACGCTGGT**

**DiR406: TCAGGTGTCGCAGGTCCAAAGGACGATGACGCTGGT**

**DiR407: AGGTGTCGCAGGTCCTTCCGACGATGACGCTGGTGC**

**DiR408: AGGTGTCGCAGGTCCTTGTGACGATGACGCTGGTGC**

**DiR409: AGGTGTCGCAGGTCCTTTAGACGATGACGCTGGTGC**

**DiR410: GTGTCGCAGGTCCTTAGCCCGATGACGCTGGTGCAC**

**DiR411: GTGTCGCAGGTCCTTAGTGCGATGACGCTGGTGCAC**

**DiR412: GTGTCGCAGGTCCTTAGATCGATGACGCTGGTGCAC**

**DiR413: GTCGCAGGTCCTTAGGAGAATGACGCTGGTGCACTT**

**DiR414: GTCGCAGGTCCTTAGGATCATGACGCTGGTGCACTT**

**DiR415: GTCGCAGGTCCTTAGGAATATGACGCTGGTGCACTT**

**DiR416: CGCAGGTCCTTAGGACGCCGACGCTGGTGCACTTCC**

**DiR417: CGCAGGTCCTTAGGACGGAGACGCTGGTGCACTTCC**

**DiR418: CGCAGGTCCTTAGGACGTGGACGCTGGTGCACTTCC**

**DiR419: CAGGTCCTTAGGACGATCCCGCTGGTGCACTTCCTG**

**DiR420: CAGGTCCTTAGGACGATTGCGCTGGTGCACTTCCTG**

**DiR421: CAGGTCCTTAGGACGATATCGCTGGTGCACTTCCTG**

**DiR422: GGTCCTTAGGACGATGAGACTGGTGCACTTCCTGAC**

**DiR423: GGTCCTTAGGACGATGATCCTGGTGCACTTCCTGAC**

**DiR424: GGTCCTTAGGACGATGAATCTGGTGCACTTCCTGAC**

**DiR425: TCCTTAGGACGATGACGGGGGTGCACTTCCTGACGT**

**DiR426: TCCTTAGGACGATGACGTAGGTGCACTTCCTGACGT**

**DiR427: TCCTTAGGACGATGACGACGGTGCACTTCCTGACGT**

**DiR428: CTTAGGACGATGACGCTCCTGCACTTCCTGACGTCG**

**DiR429: CTTAGGACGATGACGCTTATGCACTTCCTGACGTCG**

**DiR430: CTTAGGACGATGACGCTATTGCACTTCCTGACGTCG**

**DiR431: TAGGACGATGACGCTGGCCCACTTCCTGACGTCGTT**

**DiR432: TAGGACGATGACGCTGGGTCACTTCCTGACGTCGTT**

**DiR433: TAGGACGATGACGCTGGAACACTTCCTGACGTCGTT**

**DiR434: GGACGATGACGCTGGTGGTCTTCCTGACGTCGTTGT**

**DiR435: GGACGATGACGCTGGTGTCCTTCCTGACGTCGTTGT**

**DiR436: GGACGATGACGCTGGTGAGCTTCCTGACGTCGTTGT**

**DiR437: ACGATGACGCTGGTGCAGGTCCTGACGTCGTTGTCG**

**DiR438: ACGATGACGCTGGTGCATATCCTGACGTCGTTGTCG**

**DiR439: ACGATGACGCTGGTGCAACTCCTGACGTCGTTGTCG**

**DiR440: GATGACGCTGGTGCACTCACTGACGTCGTTGTCGAT**

**DiR441: GATGACGCTGGTGCACTGTCTGACGTCGTTGTCGAT**

**DiR442: GATGACGCTGGTGCACTAGCTGACGTCGTTGTCGAT**

**DiR443: TGACGCTGGTGCACTTCGGGACGTCGTTGTCGATCT**

**DiR444: TGACGCTGGTGCACTTCTAGACGTCGTTGTCGATCT**

**DiR445: TGACGCTGGTGCACTTCACGACGTCGTTGTCGATCT**

**DiR446: ACGCTGGTGCACTTCCTCCCGTCGTTGTCGATCTGG**

**DiR447: ACGCTGGTGCACTTCCTTGCGTCGTTGTCGATCTGG**

**DiR448: ACGCTGGTGCACTTCCTATCGTCGTTGTCGATCTGG**

**DiR449: GCTGGTGCACTTCCTGAGATCGTTGTCGATCTGGAG**

**DiR450: GCTGGTGCACTTCCTGATCTCGTTGTCGATCTGGAG**

**DiR451: GCTGGTGCACTTCCTGAATTCGTTGTCGATCTGGAG**

**DiR452: TGGTGCACTTCCTGACGCAGTTGTCGATCTGGAGCA**

**DiR453: TGGTGCACTTCCTGACGGTGTTGTCGATCTGGAGCA**

**DiR454: TGGTGCACTTCCTGACGAGGTTGTCGATCTGGAGCA**

**DiR455: GTGCACTTCCTGACGTCCCTGTCGATCTGGAGCACG**

**DiR456: GTGCACTTCCTGACGTCTGTGTCGATCTGGAGCACG**

**DiR457: GTGCACTTCCTGACGTCAATGTCGATCTGGAGCACG**

**DiR458: GCACTTCCTGACGTCGTCCTCGATCTGGAGCACGGA**

**DiR459: GCACTTCCTGACGTCGTGTTCGATCTGGAGCACGGA**

**DiR460: GCACTTCCTGACGTCGTAATCGATCTGGAGCACGGA**

**DiR461: ACTTCCTGACGTCGTTGCAGATCTGGAGCACGGATA**

**DiR462: ACTTCCTGACGTCGTTGGTGATCTGGAGCACGGATA**

**DiR463: ACTTCCTGACGTCGTTGAGGATCTGGAGCACGGATA**

**DiR464: CTTCCTGACGTCGTTGTCCCTCTGGAGCACGGATAGAC**

**DiR465: CTTCCTGACGTCGTTGTCTGTCTGGAGCACGGATAGAC**

**DiR466: CTTCCTGACGTCGTTGTCATTCTGGAGCACGGATAGAC**

**DiR467: CCTGACGTCGTTGTCGAGGTGGAGCACGGATAGACG**

**DiR468: CCTGACGTCGTTGTCGACATGGAGCACGGATAGACG**

**DiR469: CCTGACGTCGTTGTCGAATTGGAGCACGGATAGACG**

**DiR470: TGACGTCGTTGTCGATCACGAGCACGGATAGACGAT**

**DiR471: TGACGTCGTTGTCGATCGAGAGCACGGATAGACGAT**

**DiR472: TGACGTCGTTGTCGATCCTGAGCACGGATAGACGAT**

**DiR473: ACGTCGTTGTCGATCTGTGGCACGGATAGACGATGA**

**DiR474: ACGTCGTTGTCGATCTGACGCACGGATAGACGATGA**

**DiR475: ACGTCGTTGTCGATCTGCTGCACGGATAGACGATGA**

**DiR476: GTCGTTGTCGATCTGGAAGACGGATAGACGATGACG**

**DiR477: GTCGTTGTCGATCTGGATAACGGATAGACGATGACG**

**DiR478: GTCGTTGTCGATCTGGACTACGGATAGACGATGACG**

**DiR479: CGTTGTCGATCTGGAGCGGGGATAGACGATGACGTA**

**DiR480: CGTTGTCGATCTGGAGCCAGGATAGACGATGACGTA**

**DiR481: CGTTGTCGATCTGGAGCTTGGATAGACGATGACGTA**

**DiR482: TTGTCGATCTGGAGCACACATAGACGATGACGTAGTG**

**DiR483: TTGTCGATCTGGAGCACCAATAGACGATGACGTAGTG**

**DiR484: TTGTCGATCTGGAGCACTTATAGACGATGACGTAGTG**

**DiR485: GTCGATCTGGAGCACGGGGAGACGATGACGTAGTGA**

**DiR486: GTCGATCTGGAGCACGGTCAGACGATGACGTAGTGA**

**DiR487: GTCGATCTGGAGCACGGCAAGACGATGACGTAGTGA**

**DiR488: CGATCTGGAGCACGGATCCACGATGACGTAGTGAGA**

**DiR489: CGATCTGGAGCACGGATTAACGATGACGTAGTGAGA**

**DiR490: CGATCTGGAGCACGGATGTACGATGACGTAGTGAGA**

**DiR491: ATCTGGAGCACGGATAGGGGATGACGTAGTGAGAGA**

**DiR492: ATCTGGAGCACGGATAGCAGATGACGTAGTGAGAGA**

**DiR493: ATCTGGAGCACGGATAGTTGATGACGTAGTGAGAGATC**

**DiR494: CTGGAGCACGGATAGACTGTGACGTAGTGAGAGATC**

**DiR495: CTGGAGCACGGATAGACACTGACGTAGTGAGAGATC**

**DiR496: CTGGAGCACGGATAGACCTTGACGTAGTGAGAGATC**

**DiR497: GGAGCACGGATAGACGAACACGTAGTGAGAGATCGT**

**DiR498: GGAGCACGGATAGACGAGAACGTAGTGAGAGATCGT**

**DiR499: GGAGCACGGATAGACGACTACGTAGTGAGAGATCGT**

**DiR500: AGCACGGATAGACGATGGGGTAGTGAGAGATCGTGG**

**DiR501: AGCACGGATAGACGATGCAGTAGTGAGAGATCGTGG**

**DiR502: AGCACGGATAGACGATGTTGTAGTGAGAGATCGTGG**

**DiR503: CACGGATAGACGATGACAGAGTGAGAGATCGTGGCT**

**DiR504: CACGGATAGACGATGACCCAGTGAGAGATCGTGGCT**

**DiR505: CACGGATAGACGATGACTAAGTGAGAGATCGTGGCT**

**DiR506: CGGATAGACGATGACGTCCTGAGAGATCGTGGCTGA**

**DiR507: CGGATAGACGATGACGTTATGAGAGATCGTGGCTGA**

**DiR508: CGGATAGACGATGACGTGTTGAGAGATCGTGGCTGA**

**DiR509: GATAGACGATGACGTAGACAGAGATCGTGGCTGACG**

**DiR510: GATAGACGATGACGTAGGAAGAGATCGTGGCTGACG**

**DiR511: GATAGACGATGACGTAGCTAGAGATCGTGGCTGACG**

**DiR512: TAGACGATGACGTAGTGCCAGATCGTGGCTGACGTC**

**DiR513: TAGACGATGACGTAGTGTAAGATCGTGGCTGACGTC**

**DiR514: TAGACGATGACGTAGTGGTAGATCGTGGCTGACGTC**

**DiR515: GACGATGACGTAGTGAGCCATCGTGGCTGACGTCGC**

**DiR516: GACGATGACGTAGTGAGTAATCGTGGCTGACGTCGC**

**DiR517: GACGATGACGTAGTGAGGTATCGTGGCTGACGTCGC**

**DiR518: CGATGACGTAGTGAGAGGGCGTGGCTGACGTCGCCA**

**DiR519: CGATGACGTAGTGAGAGTCCGTGGCTGACGTCGCCA**

**DiR520: CGATGACGTAGTGAGAGCACGTGGCTGACGTCGCCA**

**DiR521: ATGACGTAGTGAGAGATTCTGGCTGACGTCGCCAGT**

**DiR522: ATGACGTAGTGAGAGATGATGGCTGACGTCGCCAGT**

**DiR523: ATGACGTAGTGAGAGATATTGGCTGACGTCGCCAGT**

**DiR524: GACGTAGTGAGAGATCGACGCTGACGTCGCCAGTCG**

**DiR525: GACGTAGTGAGAGATCGGAGCTGACGTCGCCAGTCG**

**DiR526: GACGTAGTGAGAGATCGCTGCTGACGTCGCCAGTCG**

**DiR527: CGTAGTGAGAGATCGTGAGTGACGTCGCCAGTCGAG**

**DiR528: CGTAGTGAGAGATCGTGTATGACGTCGCCAGTCGAG**

**DiR529: CGTAGTGAGAGATCGTGCTTGACGTCGCCAGTCGAG**

**DiR530: TAGTGAGAGATCGTGGCACACGTCGCCAGTCGAGTG**

**DiR531: TAGTGAGAGATCGTGGCGAACGTCGCCAGTCGAGTG**

**DiR532: TAGTGAGAGATCGTGGCCTACGTCGCCAGTCGAGTG**

**DiR533: GTGAGAGATCGTGGCTGGGGTCGCCAGTCGAGTGAC**

**DiR534: GTGAGAGATCGTGGCTGCAGTCGCCAGTCGAGTGAC**

**DiR535: GTGAGAGATCGTGGCTGTTGTCGCCAGTCGAGTGAC**

**DiR536: GAGAGATCGTGGCTGACAGCGCCAGTCGAGTGACAA**

**DiR537: GAGAGATCGTGGCTGACCCCGCCAGTCGAGTGACAA**

**DiR538: GAGAGATCGTGGCTGACTACGCCAGTCGAGTGACAA**

**DiR539: GAGATCGTGGCTGACGTTCCCAGTCGAGTGACAACT**

**DiR540: GAGATCGTGGCTGACGTGACCAGTCGAGTGACAACT**

**DiR541: GAGATCGTGGCTGACGTATCCAGTCGAGTGACAACT**

**DiR542: GATCGTGGCTGACGTCGGGAGTCGAGTGACAACTGC**

**DiR543: GATCGTGGCTGACGTCGTAAGTCGAGTGACAACTGC**

**DiR544: GATCGTGGCTGACGTCGATAGTCGAGTGACAACTGC**

**DiR545: TCGTGGCTGACGTCGCCCCTCGAGTGACAACTGCGA**

**DiR546: TCGTGGCTGACGTCGCCTATCGAGTGACAACTGCGA**

**DiR547: TCGTGGCTGACGTCGCCGTTCGAGTGACAACTGCGA**

**DiR548: GTGGCTGACGTCGCCAGGGGAGTGACAACTGCGACT**

**DiR549: GTGGCTGACGTCGCCAGCAGAGTGACAACTGCGACT**

**DiR550: GTGGCTGACGTCGCCAGATGAGTGACAACTGCGACT**

**DiR551: GGCTGACGTCGCCAGTCTGGTGACAACTGCGACTCA**

**DiR552: GGCTGACGTCGCCAGTCACGTGACAACTGCGACTCA**

**DiR553: GGCTGACGTCGCCAGTCCTGTGACAACTGCGACTCA**

**DiR554: CTGACGTCGCCAGTCGAAGGACAACTGCGACTCAGT**

**DiR555: CTGACGTCGCCAGTCGACCGACAACTGCGACTCAGT**

**DiR556: CTGACGTCGCCAGTCGATAGACAACTGCGACTCAGT**

**DiR557: GACGTCGCCAGTCGAGTTGCAACTGCGACTCAGTTG**

**DiR558: GACGTCGCCAGTCGAGTACCAACTGCGACTCAGTTG**

**DiR559: GACGTCGCCAGTCGAGTCTCAACTGCGACTCAGTTG**

**DiR560: CGTCGCCAGTCGAGTGAGGACTGCGACTCAGTTGCG**

**DiR561: CGTCGCCAGTCGAGTGAACACTGCGACTCAGTTGCG**

**DiR562: CGTCGCCAGTCGAGTGATTACTGCGACTCAGTTGCG**

**DiR563: TCGCCAGTCGAGTGACAGGTGCGACTCAGTTGCGCT**

**DiR564: TCGCCAGTCGAGTGACACATGCGACTCAGTTGCGCT**

**DiR565: TCGCCAGTCGAGTGACATTTGCGACTCAGTTGCGCT**

**DiR566: GCCAGTCGAGTGACAACACCGACTCAGTTGCGCTGA**

**DiR567: GCCAGTCGAGTGACAACGACGACTCAGTTGCGCTGA**

**DiR568: GCCAGTCGAGTGACAACCTCGACTCAGTTGCGCTGA**

**DiR569: CAGTCGAGTGACAACTGTCACTCAGTTGCGCTGAGG**

**DiR570: CAGTCGAGTGACAACTGGAACTCAGTTGCGCTGAGG**

**DiR571: CAGTCGAGTGACAACTGATACTCAGTTGCGCTGAGG**

**DiR572: GTCGAGTGACAACTGCGGGTCAGTTGCGCTGAGGAG**

**DiR573: GTCGAGTGACAACTGCGCATCAGTTGCGCTGAGGAG**

**DiR574: GTCGAGTGACAACTGCGTTTCAGTTGCGCTGAGGAG**

**DiR575: CGAGTGACAACTGCGACGGAGTTGCGCTGAGGAGTT**

**DiR576: CGAGTGACAACTGCGACCAAGTTGCGCTGAGGAGTT**

**DiR577: CGAGTGACAACTGCGACATAGTTGCGCTGAGGAGTT**

**DiR578: AGTGACAACTGCGACTCCCTTGCGCTGAGGAGTTGT**

**DiR579: AGTGACAACTGCGACTCTATTGCGCTGAGGAGTTGT**

**DiR580: AGTGACAACTGCGACTCGTTTGCGCTGAGGAGTTGT**

**DiR581: TGACAACTGCGACTCAGGGGCGCTGAGGAGTTGTGC**

**DiR582: TGACAACTGCGACTCAGACGCGCTGAGGAGTTGTGC**

**DiR583: TGACAACTGCGACTCAGCAGCGCTGAGGAGTTGTGC**

**DiR584: ACAACTGCGACTCAGTTAGGCTGAGGAGTTGTGCTT**

**DiR585: GACAACTGCGACTCAGTTTAGCTGAGGAGTTGTGCTT**

**DiR586: ACAACTGCGACTCAGTTCTGCTGAGGAGTTGTGCTT**

**DiR587:** **AACTGCGACTCAGTTGCAGTGAGGAGTTGTGCTTGT**

**DiR588: AACTGCGACTCAGTTGCTATGAGGAGTTGTGCTTGTG**

**DiR589: AACTGCGACTCAGTTGCCTTGAGGAGTTGTGCTTGT**

**DiR590: CTGCGACTCAGTTGCGCACAGGAGTTGTGCTTGTGG**

**DiR591: CTGCGACTCAGTTGCGCGAAGGAGTTGTGCTTGTGG**

**DiR592: CTGCGACTCAGTTGCGCCTAGGAGTTGTGCTTGTGG**

**DiR593: GCGACTCAGTTGCGCTGCCGAGTTGTGCTTGTGGAC**

**DiR594: GCGACTCAGTTGCGCTGTAGAGTTGTGCTTGTGGAC**

**DiR595: GCGACTCAGTTGCGCTGGTGAGTTGTGCTTGTGGAC**

**DiR596: GACTCAGTTGCGCTGAGTGGTTGTGCTTGTGGACGA**

**DiR597: GACTCAGTTGCGCTGAGACGTTGTGCTTGTGGACGA**

**DiR598: GACTCAGTTGCGCTGAGCTGTTGTGCTTGTGGACGA**

**DiR599: CTCAGTTGCGCTGAGGAAGTGTGCTTGTGGACGAAG**

**DiR600: CTCAGTTGCGCTGAGGACCTGTGCTTGTGGACGAAG**

**DiR601: CTCAGTTGCGCTGAGGATATGTGCTTGTGGACGAAG**

**DiR602: CAGTTGCGCTGAGGAGTACTGCTTGTGGACGAAGTA**

**DiR603: CAGTTGCGCTGAGGAGTGATGCTTGTGGACGAAGTA**

**DiR604: CAGTTGCGCTGAGGAGTCTTGCTTGTGGACGAAGTA**

**DiR605: GTTGCGCTGAGGAGTTGACCTTGTGGACGAAGTACC**

**DiR606: GTTGCGCTGAGGAGTTGGACTTGTGGACGAAGTACC**

**DiR607: GTTGCGCTGAGGAGTTGCTCTTGTGGACGAAGTACC**

**DiR608: TGCGCTGAGGAGTTGTGGGTGTGGACGAAGTACCGA**

**DiR609: TGCGCTGAGGAGTTGTGACTGTGGACGAAGTACCGA**

**DiR610: TGCGCTGAGGAGTTGTGTATGTGGACGAAGTACCGA**

**DiR611: CGCTGAGGAGTTGTGCTACTGGACGAAGTACCGACA**

**DiR612: CGCTGAGGAGTTGTGCTGATGGACGAAGTACCGACA**

**DiR613: CGCTGAGGAGTTGTGCTCTTGGACGAAGTACCGACA**

**DiR614: CTGAGGAGTTGTGCTTGACGACGAAGTACCGACAGG**

**DiR615: CTGAGGAGTTGTGCTTGGAGACGAAGTACCGACAGG**

**DiR616: CTGAGGAGTTGTGCTTGCTGACGAAGTACCGACAGG**

**DiR617: GAGGAGTTGTGCTTGTGTGCGAAGTACCGACAGGTC**

**DiR618: GAGGAGTTGTGCTTGTGACCGAAGTACCGACAGGTC**

**DiR619: GAGGAGTTGTGCTTGTGCTCGAAGTACCGACAGGTC**

**DiR620: GGAGTTGTGCTTGTGGATCAAGTACCGACAGGTCTG**

**DiR621: GGAGTTGTGCTTGTGGAGAAAGTACCGACAGGTCTG**

**DiR622: GGAGTTGTGCTTGTGGAATAAGTACCGACAGGTCTG**

**DiR623: AGTTGTGCTTGTGGACGGGGTACCGACAGGTCTGAC**

**DiR624: AGTTGTGCTTGTGGACGCCGTACCGACAGGTCTGAC**

**DiR625: AGTTGTGCTTGTGGACGTTGTACCGACAGGTCTGAC**

**DiR626: TTGTGCTTGTGGACGAAAGACCGACAGGTCTGACCG**

**DiR627: TTGTGCTTGTGGACGAACCACCGACAGGTCTGACCG**

**DiR628: TTGTGCTTGTGGACGAATAACCGACAGGTCTGACCG**

**References**
